# Nucleolar dynamics are determined by the ordered assembly of the ribosome

**DOI:** 10.1101/2023.09.26.559432

**Authors:** Jessica Sheu-Gruttadauria, Xiaowei Yan, Nico Stuurman, Riley O. Ogrean, Sabrina Lin, Stephen N. Floor, Ronald D. Vale

## Abstract

Ribosome biogenesis occurs in the nucleolus, a biomolecular condensate whose material properties are thought to be important for function. However, the molecular basis of nucleolar dynamics and their relationship to ribosome assembly remain incompletely understood. We present a platform for <u>hi</u>gh-throughput fluorescence <u>r</u>ecovery <u>a</u>fter <u>p</u>hotobleaching (HiT-FRAP) and use it to screen hundreds of genes for their impact on dynamics of the nucleolar scaffold nucleophosmin (NPM1). We find that NPM1 dynamics and nucleolar morphology are sensitive to ribosome assembly state: accumulation of early preribosomal intermediates slows NPM1 dynamics and compacts the condensate, while accumulation of abortive late precursors accelerates dynamics and disrupts condensate integrity. These opposing biophysical states correlate with the strength of NPM1-pre-ribosome interactions. Importantly, mutations in the NPM1 intrinsically disordered region that alter pre-ribosome binding directly tune nucleolar dynamics. These results establish that ribosomal precursor assembly state determines nucleolar material properties through the valency of scaffold-pre-ribosome interactions and introduce HiT-FRAP as a platform for interrogating condensate dynamics broadly.

**TOC:** Sheu-Gruttadauria et al. show that the material properties of the nucleolus are shaped by the assembly state of ribosomal precursors, which tune nucleolar dynamics and morphology through their interactions with the nucleolar scaffolding protein NPM1.

## Introduction

Biomolecular condensates are membraneless organelles that are broadly implicated in many cellular processes, including nearly every step in the life cycle of an RNA [1–4]. Over the last decade, increasing evidence suggests that many condensates form through biological phase transitions, driven by multivalent interaction networks that form between nucleic acids and intrinsically disordered regions (IDRs) in constituent proteins [5]. These interaction networks imbue condensates with an array of dynamic biophysical properties, including exchange with the environment, internal mixing, viscosity, and surface tension, resulting in material properties that range from liquid to solid-like. It is thought that these biophysical properties influence the biological function of condensates, as pathological mutations that disrupt phase transitions or promote aggregation are associated with disease [1, 4]. Emerging work has begun to uncover the molecular principles that underlie condensate dynamics in reconstituted phase separating systems [6–9]. However, mechanistically connecting these features to condensate function in living systems has remained challenging.

One of the first identified and largest biomolecular condensates in the cell is the nucleolus, whose primary function is coordinating ribosome biogenesis, a process essential for cell viability, growth, and proliferation [10–12]. Decades of studies have provided detailed molecular insight into this process [13]. In humans, ribosome biogenesis begins with transcription of the 47S pre-ribosomal RNA (rRNA) by RNA polymerase I (Pol I), which is processed together with the RNA Pol III-transcribed 5S rRNA to yield the small subunit (SSU) 18S and large subunit (LSU) 28S, 5.8S, and 5S rRNAs (Fig. S1A). This rRNA scaffold undergoes modification, processing, and folding, concomitant with recruitment of 80 ribosomal proteins, all of which is orchestrated by over 200 assembly factors [13–15]. The SSU and LSU proceed through over a dozen discrete intermediates, many of which have been structurally characterized [13, 16–18], providing a detailed molecular model for stages of ribosome assembly.

In contrast to this precise understanding of ribosome biogenesis, the molecular underpinnings of nucleolar form and how they relate to ribosome assembly are only beginning to be understood. On the mesoscale, the nucleolus is composed of three concentric layers enriched for factors involved in sequential steps of ribosome assembly: the fibrillar center (FC) where Pol I transcription occurs, the dense fibrillar component (DFC) where SSU assembly and rRNA processing factors localize, and the granular component (GC) that houses factors involved in maturation of the pre-SSU and assembly of the pre-LSU [19–21] (Fig. S1A). A key scaffolding protein of the GC is nucleophosmin (NPM1), a pentameric protein bearing acidic and basic intrinsically disordered regions that mediate multivalent interactions with ribosomal proteins, assembly factors, and rRNA, promoting phase separation into the GC [22–24]. NPM1 is one of the most abundant nucleolar proteins and one of the first identified and best studied phase-separating scaffolds in the nucleolus [22, 23, 25], making it an attractive model for understanding how condensate properties emerge from constituent interactions.

Nucleolar assembly through phase separation leads to several biophysical properties that have been proposed to facilitate ribosome biogenesis. Firstly, it is thought that the three nucleolar layers are immiscible subphases kept distinct by differences in their surface tensions and viscosities, thereby spatially organizing ribosome biogenesis in an assembly line [26]. The interfaces of these subphases may be important for molecular hand-off between stages of assembly [27, 28]. Moreover, in vitro and bioinformatic work suggests that the interaction valency of pre-ribosomes with nucleolar scaffolds decreases during maturation through folding and processing of the rRNA, binding of ribosomal proteins, and departure of assembly factors [24, 29, 30], promoting the thermodynamic release of mature subunits. Lastly, nucleolar material properties, often described as liquid-like, are thought to be essential for proper ribosome biogenesis [25]. Consistent with this idea, optogenetic clustering of nucleolar scaffolds leads to rigidification and perturbed rRNA processing [31]. Nucleolar morphology is also often disrupted in cancer, aging, and neurodegeneration [32].

Together, these observations support a compelling hypothesis that the biophysical features of the nucleolus are intrinsically coupled with its function in ribosome assembly. Indeed, recent advances have connected the multilayered organization of the nucleolus to rRNA processing, demonstrating that its concentric nature is templated by rRNA and its hierarchical maturation [28, 33, 34]. Moreover, previous imaging-based screens have coupled nucleolar morphology to defects in ribosome biogenesis [35–39]. However, what determines the dynamic material properties of the nucleolus and, importantly, how these are mechanistically tied to ribosome assembly remain far less understood. Current understanding of nucleolar dynamics come from in vitro reconstitution of isolated nucleolar constituents, as the field currently lacks tools for systematically measuring condensate dynamics in living cells. As a result, the molecular basis for nucleolar dynamics remains incompletely defined, limiting our ability to connect the emergent biophysical features of condensates to their function.

Here, we introduce a platform to systematically measure dynamics of biomolecular condensates in live cells. The most readily measurable biophysical property of condensates in living systems is macromolecular diffusivity, which is typically assessed using fluorescence recovery after photobleaching, or FRAP [40]. FRAP is a robust and quantitative technique that is widely accessible using commercial microscopes. However, it is also a low-throughput and largely manual technique. To overcome this limitation, we developed a method for <u>Hi</u>gh-<u>T</u>hroughput <u>F</u>luorescence <u>R</u>ecovery <u>A</u>fter <u>P</u>hotobleaching, which we call HiT-FRAP. This strategy allows FRAP to be used as a readout in arrayed genetic screens, enabling discovery of factors that influence condensate dynamics.

We apply HiT-FRAP to screen hundreds of genes for their impact on the dynamics of NPM1, using it as a proxy for dynamics in the GC. When combined with analysis of nucleolar morphology, we identify distinct nucleolar phenotypes associated with specific stages in ribosome assembly, as well as an unexpected connection to mRNA processing pathways. Strikingly, across phenotypic clusters, changes in NPM1 dynamics are mirrored by changes in nucleolar morphology and NPM1 partitioning, suggesting that these readouts together report on the material state of the condensate. Characterizing ribosome biogenesis across these states reveals that they correspond to changes in pre-ribosome composition within the nucleolus: accumulation of early intermediates decreases NPM1 dynamics while accumulation of abortive late LSU intermediates increases exchange. These trends correlate directly with NPM1 partitioning into the nucleolus and the strength of its interactions with ribosomal intermediates, which suggests that the interactions that drive phase separation also dictate condensate material properties. We further demonstrate that mutations in the NPM1 intrinsically disordered region that alter pre-ribosome binding strength directly tune both partitioning and dynamics. Together, these results establish that the assembly state of ribosomal precursors determines the dynamic biophysical properties of the nucleolus, providing a direct mechanistic tie between ribosome biogenesis and nucleolar material properties.

## Results

### HiT-FRAP: A platform for performing scalable automated FRAP

To identify factors that guide macromolecular dynamics in biomolecular condensates, we developed a platform to perform scalable, fully automated FRAP experiments called HiT-FRAP (Fig. 1A). This pipeline uses data-adaptive imaging strategies to automatically identify and bleach fluorescently labeled organelles in living cells and acquire movies to measure fluorescence recovery (Fig. 1B), eliminating the need for manual designation of regions of interest and increasing experimental throughput. We also developed an automated analysis pipeline that quantifies fluorescence recovery for all bleach points across all acquired positions, normalizing to non-bleached organelles. We fit recovery data to a simple exponential to prevent over-fitting and export parameters that describe dynamics (t_1/2_ and mobile fraction, which represent time to 50% recovery and percentage of mobile molecules, respectively).

**Figure 1:**
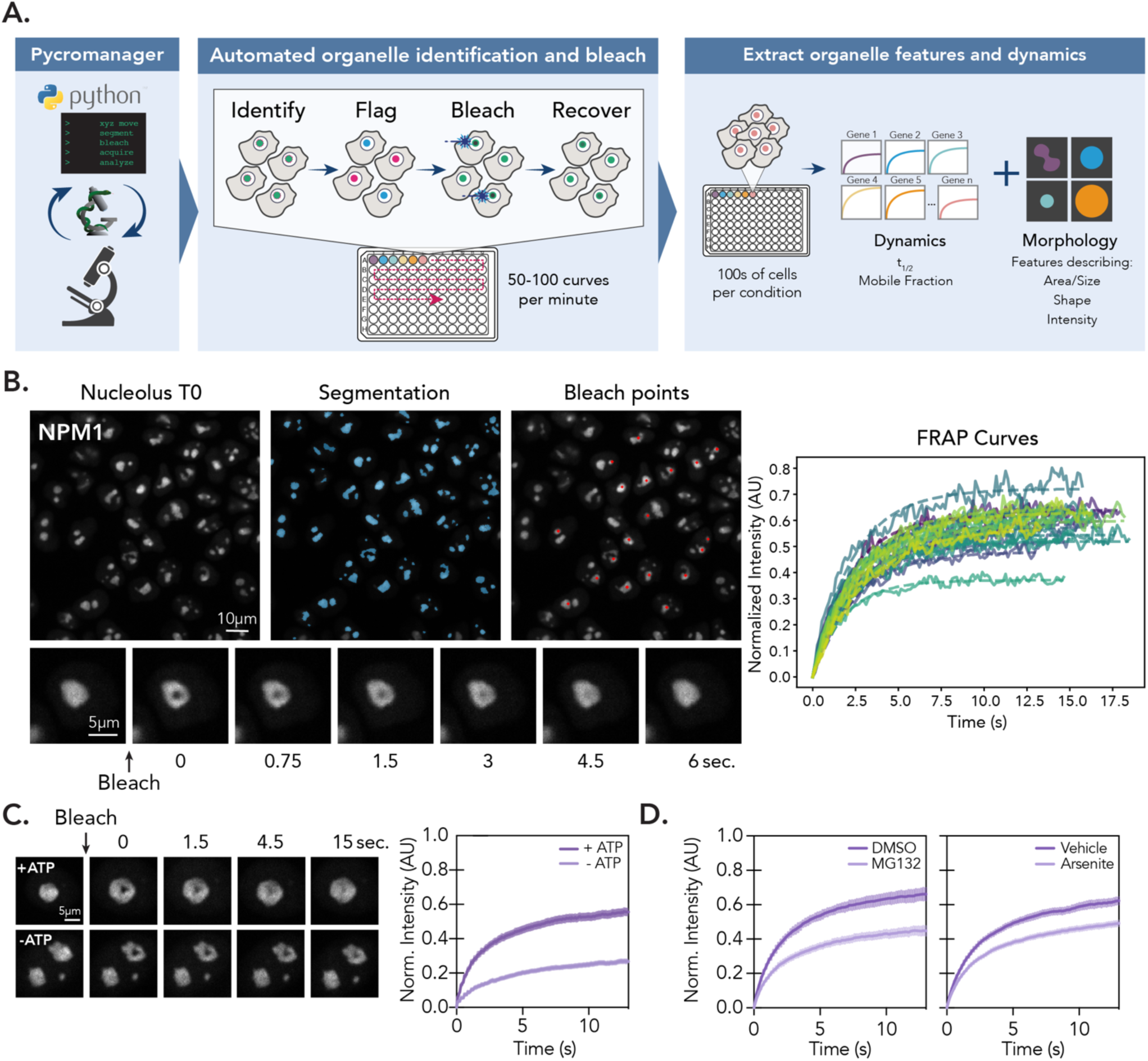
HiT-FRAP: A platform for performing scalable, automated FRAP experiments. (A) Schematic overview of HiT-FRAP pipeline. See also: Methods. (B) Example images show automated FRAP steps. Left, segmentation of fluorescently labeled nucleoli (blue masks, see methods) and bleach point determination (red points). Below, zoom of single nucleolus bleach and recovery time course. Right, normalized FRAP curves generated from a single field of view. Raw intensities shown as solid line, single-exponential curve fit shown as dotted line. (C) NPM1-mNeonGreen cell lines were treated with 10 mM sodium azide and 6 mM 2-deoxyglucose for 10 min to deplete cellular ATP prior to bleaching experiment. Left, images of single nucleolus bleach and recovery time course for vehicle and ATP deplete. Right, FRAP curves (n = 250 nucleoli, error bars are 95% CI). (D) NPM1-mNeonGreen cell lines were treated with 10 µM MG132 for 1 hr (left) or 200 µM sodium arsenite for 1 hr prior to bleaching experiment. (n = 250 nucleoli, error bars are 95% CI).

We applied HiT-FRAP to interrogate dynamics in the largest nucleolar subcompartment, the GC, using NPM1 as a readout. We chose HeLa cells as a model system in part because they lack functional p53, which is typically activated by nucleolar stress, allowing us to study disruptions in ribosome biogenesis without confounding cell death responses. To monitor NPM1 dynamics, we used CRISPR-based tagging of the endogenous NPM1 gene to generate cell lines with C-terminal fluorescent protein tags: a homozygous NPM1-mNeonGreen line and a heterozygous NPM1-mScarlet line, the latter of which maintains wild-type NPM1 expression levels (Fig. S1B). Both lines show similar NPM1 localization, exhibit FRAP recovery kinetics consistent with published values [25], and have normal rRNA processing relative to unedited cells (Fig. S1C-D), validating their use as reporters of nucleolar dynamics.

To validate HiT-FRAP, we measured relative differences in NPM1 dynamics using perturbations previously reported to impact exchange by FRAP. Firstly, we rapidly depleted ATP with 2-deoxyglucose (2-DG) and sodium azide to block both glycolysis and oxidative phosphorylation, which had been previously shown to decrease NPM1 mobility [26]. Similarly, we found that NPM1 exhibited slower recovery (∼1.5-fold) and a lower percentage of mobile molecules (∼2-fold) as compared to vehicle alone (Fig. 1C). In addition, previous reports have shown that inhibition of the proteasome disrupts nucleolar assembly features, in keeping with its proposed role in proteostasis [41–43]. Consistent with these observations, we found that exposure to the proteasomal inhibitor MG132 decreased NPM1 mobility by ∼1.4-fold (Fig. 1D). Lastly, recent work has demonstrated that cellular stress can lead to rigidification of the nucleolus [44]. Therefore, we treated our cells with sodium arsenite and found a ∼1.3 fold decrease in mobile fraction (Fig. 1D), consistent with reported values. Together, these results demonstrate that HiT-FRAP enables automated FRAP measurements of NPM1 that robustly report on perturbations to dynamics.

### Systematic identification of factors that impact NPM1 dynamics

We next used HiT-FRAP to systematically identify factors that regulate NPM1 dynamics in the nucleolus. We focused initially on RNA helicases, which have been proposed to modulate condensate dynamics through their effects on RNA-driven phase separation [45–48]. Many RNA helicases also localize to the nucleolus where they help coordinate ribosome biogenesis [49]. Using an arrayed siRNA library, we depleted 65 RNA helicases and 9 phylogenetically related helicase pairs (Fig. S2A) in our mNeonGreen-tagged cell line and monitored NPM1 dynamics by HiT-FRAP. We identified 13 RNA helicases or helicase pairs whose depletion significantly affected t_1/2_, and 9 that altered mobile fraction (Fig. 2A, Fig. S2B). Hits were validated using de-pooled siRNAs and confirmed by western blot for select targets (Fig. S2C and D) and were highly reproducible in the mScarlet cell line (Fig. S2E). Because mobile fraction was less reproducible across replicates and cell lines, however, we focused subsequent analysis on factors affecting exchange kinetics.

**Figure 2:**
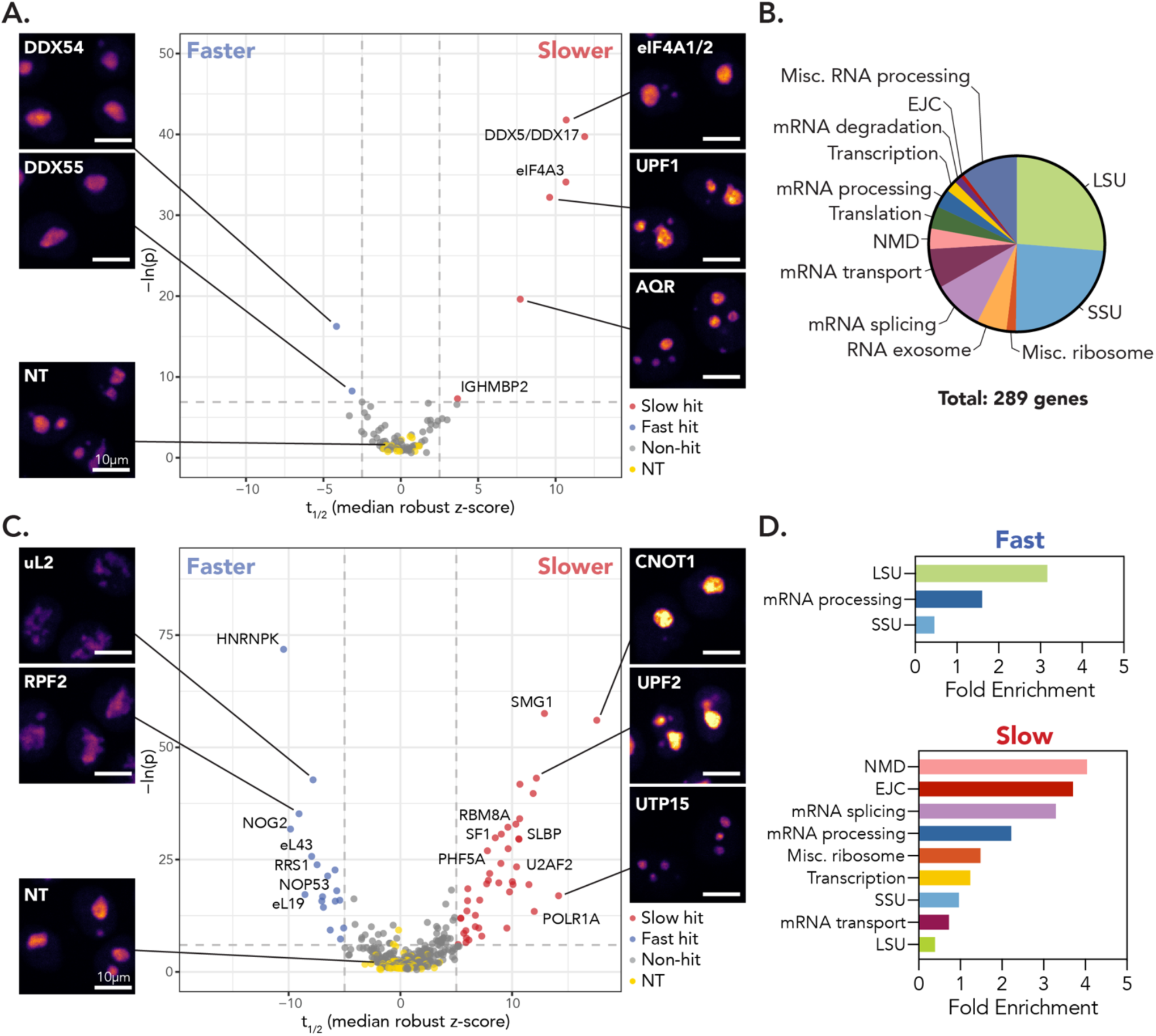
HiT-FRAP identifies RNA helicases and RNA processing factors that impact nucleolar dynamics. (A) Selected images and volcano plot for t_1/2_ for primary RNA helicase screen, highlighting helicase hits that resulted in increased (red, “slower”) or decreased (blue, “faster”) t_1/2_ relative to non-targeting control cells (yellow; FDR < 0.05 and z-cutoff of ± 2.5 shown as dotted lines, see Methods). Non-significant gene targets shown in gray. x-axis plots robust z-score for t_1/2_ (see Methods). Images are scaled equally and colored linearly with mp-inferno LUT to show NPM1 intensity. Scale bar is 10 μm. (B) Functional gene distribution of secondary screen by manual annotation. (C) Selected images and volcano plot as in (A) for secondary screen. (FDR < 0.05 and z-cutoff of ± 4, shown as dotted lines). (D) Gene enrichment for functional groups, determined by manual annotation, for “fast” (smaller t_1/2_) and “slow” (larger t_1/2_) hits.

Of the 13 helicases influencing t_1/2_, 8 are nucleolar or implicated in ribosome biogenesis (Fig. 2A). Helicases whose depletion accelerated recovery (DDX18 [50], DDX54 [51], DDX55 [52]) are involved in LSU assembly, while three of four helicases whose depletion slowed exchange (DDX41 [53], eIF4A3 [54] and IGHMBP2 [55]) coordinate early rRNA processing through the SSU processome. The exception was co-depletion of DDX5 and DDX17, which remodel the LSU 28S rRNA [56]. Notably, these and other “slow” hits (AQR, DHX8, DHX38, eIF4A1+eIF4A2, UPF1) also function in pre-mRNA processing, NMD, and translation [57], complicating interpretation of their nucleolar phenotypes.

Therefore, to distinguish direct from indirect effects, we designed a secondary screen of 290 gene depletions (Fig. 2B), comprising ribosomal proteins and assembly factors (51%) alongside STRING-database interactors of the helicase hits spanning diverse RNA processing pathways [58]. This screen yielded 55 “slow” and 21 “fast” hits (Fig. 2C). Consistent with the primary screen, “fast” hits were enriched for LSU assembly factors and ribosomal proteins, suggesting that accelerated NPM1 exchange reflects broad disruption of LSU maturation (Fig. 2D). In contrast, “slow” hits were enriched for mRNA splicing, processing, and NMD pathways, which suggest that DDX5+DDX17, DDX41, eIF4A3, and IGHMBP2 likely influence nucleolar dynamics downstream of their non-ribosome biogenesis roles. Together, these screens reveal that NPM1 exchange is accelerated by disruption of LSU assembly and slowed by disruption of mRNA processing pathways.

### Phenotypic clustering reveals four nucleolar biophysical states associated with distinct biological processes

Many perturbations that affected NPM1 dynamics also produced notable changes in nucleolar morphology (Fig. 2A and C), consistent with both reporting on the condensate material state. We therefore reasoned that integrating these readouts may better distinguish biologically meaningful phenotypic classes than dynamics alone. We expanded our analysis to include morphological descriptors (mean intensity, area, circularity, and eccentricity) alongside our dynamic measures (t_1/2_ and mobile fraction, Fig. 3A) and integrated these features using principal component analysis (PCA) to visualize phenotypic relationships across the dataset (Fig. 3A-C, Fig. S3A). Phenotype scores used in PCA incorporated both statistical significance and effect size relative to non-targeting controls to minimize false positive and batch effects (see Methods). Hits were defined by Euclidean distance from the NT cluster (top 15% cutoff) and reproducibility in at least two of three biological replicates (Fig. S3B). Phenotypes were largely consistent in mScarlet-tagged lines (Fig. S3C), and major morphological features were confirmed using orthogonal GC and DFC markers (S4A-D). Phenotypes emerged within 24-48 hours of gene depletion, which suggests that they do not result from long-term growth defects (Fig. S4E).

**Figure 3:**
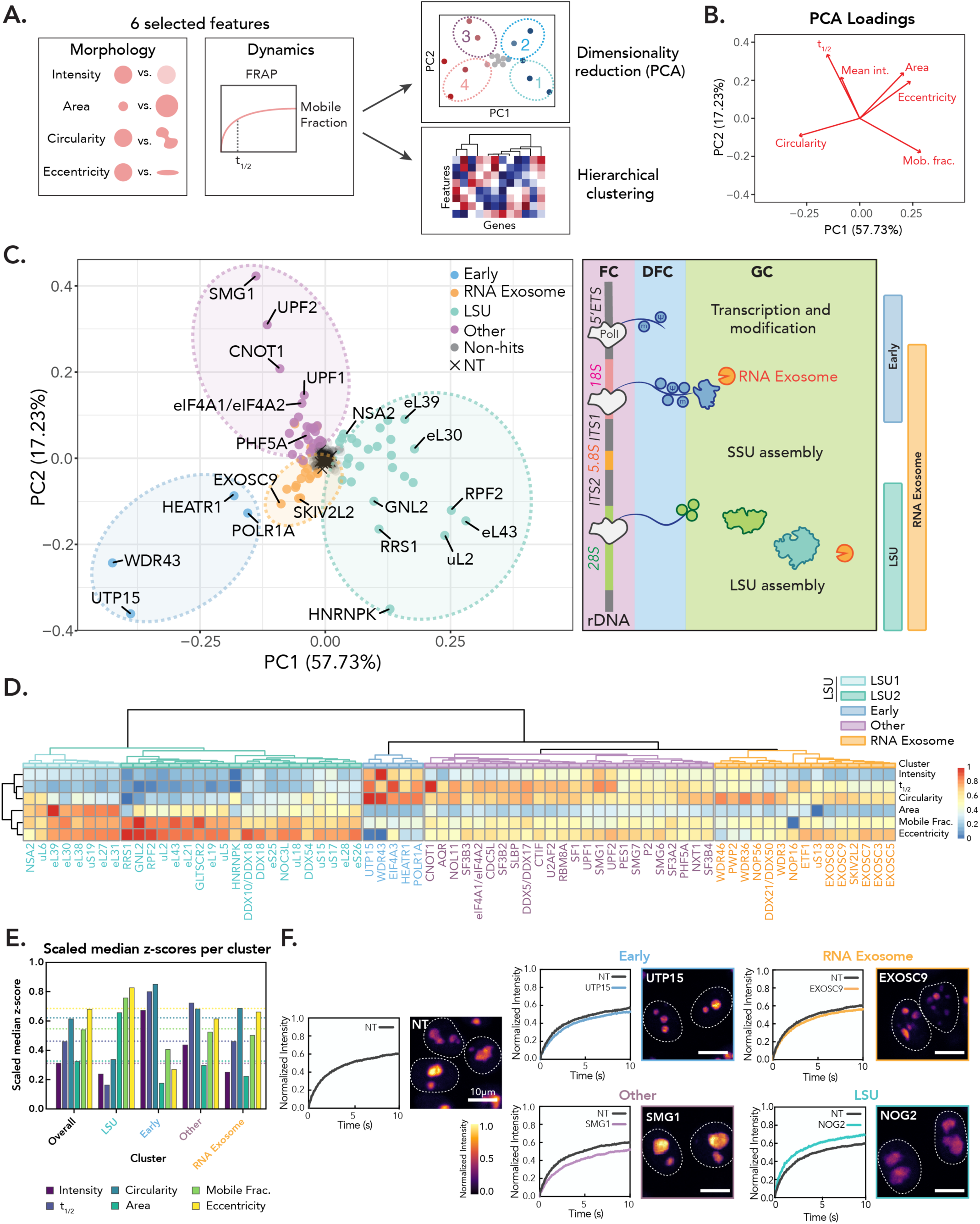
Phenotype clustering identifies gene groups associated with specific nucleolar signatures. (A) Schematic overview of phenotypic clustering analysis (see also: Methods). (B) PCA loadings for each nucleolar feature. (C) PCA colored by phenotype cluster. Schematic of nucleolar ribosome biogenesis stages shown on right, with phenotypic classes associated with stages indicated. (D) Hierarchical clustering analysis for hits (see Methods) using robust z-scores. Z-scores were scaled from 0 to 1. Clusters colored by phenotypic group. (E) Median scaled z-scores within phenotypic clusters for measured features. Dotted line shows median values for all hits that went into the analysis. (F) Representative images and FRAP curve for select hit in each phenotypic cluster. Images are scaled equally and colored with mpl-inferno LUT to show relative intensity differences in NPM1. White dotted line indicates nuclear boundary. Scale bar is 10 μm. FRAP curve of NT control and gene depletion shown (n = 250 nucleoli, error bars are 95% CI).

Grouping hits by function revealed that genes with related functions clustered together in PCA space, forming four distinct phenotypic groupings, which we term nucleolar biophysical states (Fig. 3C and D, Fig. S4F). These groupings were validated by hierarchical clustering of phenotypic z-scores, which recovered similar gene clusters (Fig. 3D). Three of the four states were associated with ribosome biogenesis: early rRNA processing and transcription factors (“Early,” Fig. 3D-F), the RNA exosome, which coordinates rRNA end-processing throughout biogenesis (“RNA Exosome,” Fig. 3D-F), and factors involved in LSU assembly (“LSU,” Fig. 3D-F). Clustering analysis further subdivided the LSU group into two related sub-clusters (Fig. 3D, see Supplementary Text and Figure S5 for further discussion of these groups). The fourth state (“Other,” Fig. 3D) comprised factors with no known role in ribosome biogenesis and included most “slow” hits identified by HiT-FRAP. Notably, both PCA and hierarchical clustering of features revealed that certain dynamic and morphological features co-vary across the dataset, with faster NPM1 dynamics tracking with decreased NPM1 intensity, increased nucleolar size, and increased irregularity, and slower dynamics tracking with increased circularity and increased NPM1 intensity (Fig. 3B and D). This coherent covariance suggests that dynamics and morphology together report on the material state of the nucleolar condensate. Collectively, these analyses identify four nucleolar biophysical states that each reflect distinct biological processes, in particular stages of ribosome assembly.

### Nucleolar phenotypes reflect disruptions at distinct stages of ribosome biogenesis

Two of the four phenotypic states (“Early” and “RNA Exosome”) were enriched for factors involved in rRNA transcription or early rRNA processing. Consistent with previous reports [59], depletion of both groups caused nucleolar shrinking and decreased NPM1 dynamics, with RNA Exosome depletions showing greater morphological irregularity (Fig. 3C-F). This fragmented morphology was recapitulated by acute treatment with Pol I inhibitors ActD (0.04 µg/mL [60, 61]) and CX5461 [62] (500 nM), confirming that these phenotypes reflect decreased rRNA transcription (Fig. S6A). Consistent with a role for pre-ribosome intermediates in maintaining nucleolar architecture, rRNA FISH using probes against the 5′ETS, ITS1, and ITS2 (removed during early/SSU-, SSU-, and LSU-associated maturation stages, respectively) revealed depletion of all rRNA precursors from the nucleolus under both inhibitor treatment and knockdown of the core RNA exosome helicase SKIV2L2, with Northern blot confirming distinct processing defects for each condition (Fig. 4A, S6B, S7).

**Figure 4:**
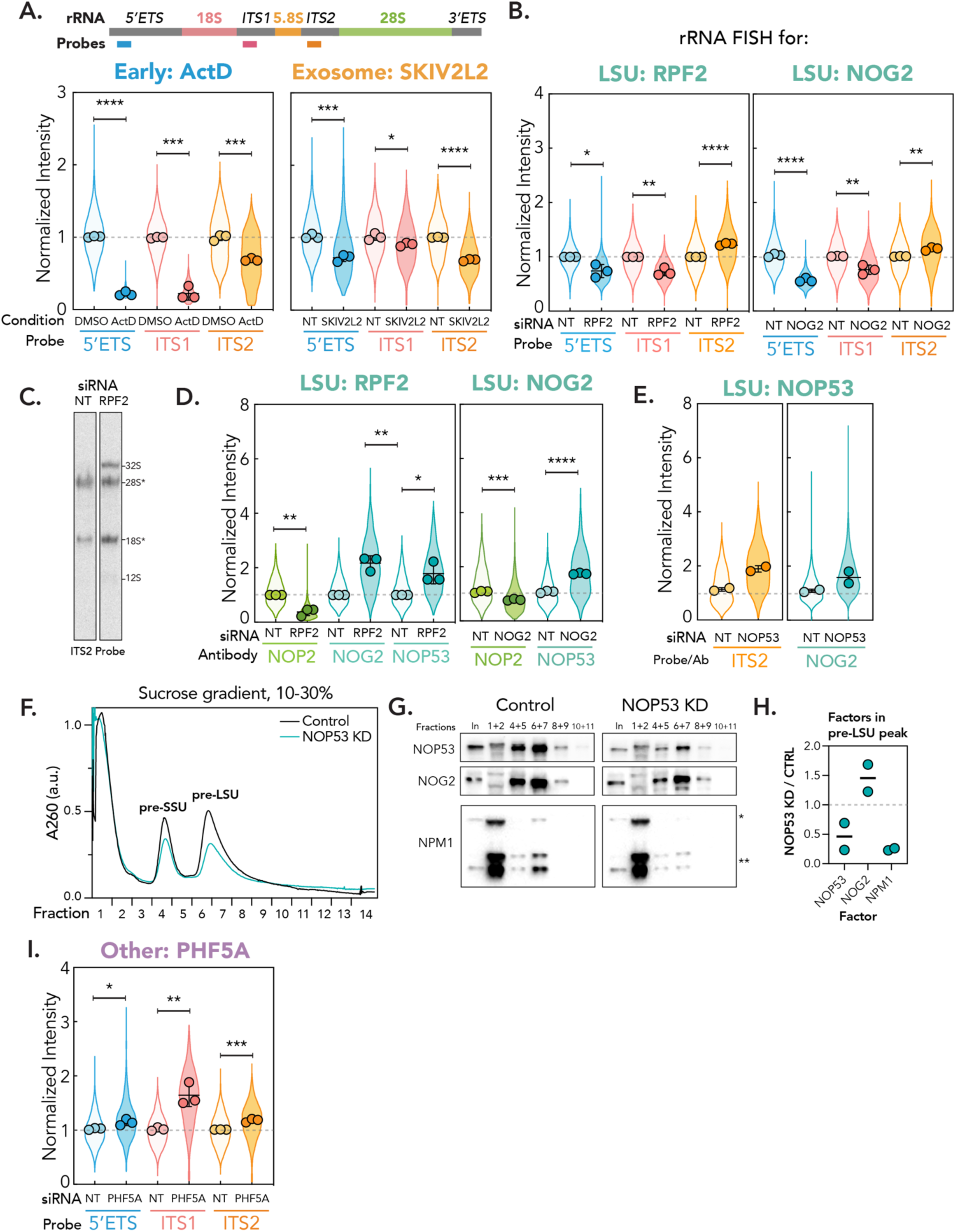
Distinct nucleolar phenotypes are associated with discrete stages in ribosome biogenesis. (A) rRNA FISH for Early (ActD) and Exosome cluster (knockdown of SKIV2L2). Schematic shows position of probes in 47S rRNA. Violin plots show spread of individual data points across three biological replicates (n = 600 nucleoli per replicate, points represent means of replicates, error bars are SD). The dotted line shows non-targeting control level. p-values calculated using two-tailed unpaired t-test between biological replicates. * p < 0.05, ** p<0.01, *** p < 0.001, **** p<0.0001. (B) rRNA FISH for representative hits RPF2 and NOG2 in LSU cluster, as in (B). (C) ITS2 northern blot for NT and RPF2-knockdown cells. 32S indicated, non-specific 28S and 18S bands shown. Gel has been cropped to remove lanes for clarity, full gel available in supplement. (D) Nucleolar immunofluorescence of LSU intermediate markers for LSU representative hits RPF2 and NOG2. Three biological replicates were performed (n = 600 nucleoli per replicate, spread of individual nucleoli shown as violin, error bars are SD). Dotted line shows non-targeting control level. p-values calculated using two-tailed unpaired t-test between biological replicates. n.s. = not significant, * p < 0.05, ** p<0.01, *** p < 0.001. (E) Left, ITS2 FISH for representative LSU hit NOP53. Right, nucleolar immunofluorescence for LSU intermediate marker NOG2 for representative LSU hit NOP53. (Two biological replicates shown, n = 600 nucleoli per replicate, spread of individual nucleoli shown as violin, error bars are SD). (F) Sucrose gradient (10-30% w/v) fractionation of nuclear pre-ribosomal complexes isolated from HeLa cells transfected with NOP53 or with control siRNA. (G) Western blot showing NOP53, NOG2, and NPM1 from pooled fractions as indicated. * = NPM1-mScarlet, ** = untagged NPM1. (H) Quantification of LSU intermediate assembly factors from western blot that co-sediment with pre-LSU peak. Normalized to the area under the curve for pooled fractions 6-9, then normalized to control. Two replicates shown, bar is mean. (I) rRNA FISH for representative gene hit PHF5A in “Other” phenotypic cluster, as in (B). n.s. = not significant, * p < 0.05, ** p<0.01, *** p < 0.001.

The largest group of genes we identified were involved in LSU maturation, in keeping with the GC’s primary role in LSU assembly (“LSU”). The depletion of these genes leads to an increase in nucleolar size and irregularity and a substantial decrease in NPM1 t_1/2_, which suggests disruption of condensate integrity and increased dynamics (Fig. 3C-F). This morphological phenotype is consistent with previous nucleolar screens that reported similar increases in nucleolar size and irregularity upon perturbation of “LSU”-associated ribosomal proteins and assembly factors [35, 38, 39]. Moreover, we also find that disruption of “LSU” genes in p53-proficient, hTERT-RPE1 cells has a similar impact on nucleolar morphology (Fig. S6C and Supplementary Text), which suggests that this phenotypic cluster might represent a general nucleolar response to LSU disruption.

When mapping the factors we identify onto pre-LSU structures, we find that they are primarily located around late-forming structural landmarks including the inter-subunit interface, L1 stalk, and the binding site for the central protuberance (CP; Fig. S5B and C) whose perturbation causes accumulation of late-stage abortive intermediates in yeast [63–65]. We therefore hypothesized the nucleolar changes we observe might result from accumulation of abortive intermediates in the nucleolus. In support of this hypothesis, knockdown of representative “LSU” hits RPF2 (coordinates 5S RNP incorporation and CP maturation) and NOG2 (a GTPase involved in late pre-LSU maturation) led to a significant nucleolar depletion of the 5′ETS and ITS1 and accumulation of ITS2 in the nucleolus (Fig. 4B, S8). Northern blotting confirmed an accumulation of the 32S and 12S pre-rRNAs upon depletion of RPF2 as well as “LSU” hit NOP53 (responsible for RNA exosome recruitment for ITS2 processing), suggesting that while ITS2 can be cleaved, end-processing is impaired (Fig. 4C and S6B). As ITS2 is processed immediately after nucleolar export, these data are consistent with a pre-export assembly stall, accumulation of abortive intermediates, and depletion of mature LSUs, as supported by reduced mature 28S levels by qPCR (Fig. S8A).

To further characterize these stalled intermediates, we turned to recent structural work that identified at least eight distinct nucleolar LSU intermediates, termed States A-H [16]. Many of the factors that we identified in our “LSU” phenotype group are associated with key maturation events in transitions between late States F through H (Fig. S5B and C), which suggests that disruption of these factors might result in nucleolar accumulation of stalled complexes that resemble these states. To explore this idea, we determined the nucleolar localization of factors associated with States F through H by immunofluorescence: NOP2, an rRNA methyltransferase associated with pre-LSU states C-F, and NOP53 and NOG2, which both bind to late intermediates G and H. When we depleted either RPF2 and NOG2, we observed a strong depletion of NOP2 from the nucleolus and enrichment for NOP53 and NOG2 (Fig. 4D and S8B). To confirm these results in an additional LSU gene, we also depleted NOP53 and saw a similar nucleolar accumulation of both ITS2 and NOG2 (Fig. 4E). To determine if these factors are part of developing pre-LSUs, we performed sucrose gradient sedimentation of nuclear pre-ribosomes in NOP53 knockdown cells. We found that NOG2 is not enriched in soluble fractions and instead co-sediments with pre-LSUs, confirming that the IF factors we observe are part of developing intermediates (Fig. 4G and H). Together with the accumulation of the 32S and nucleolar ITS2, these results suggest that when LSU group factors are depleted, very late assembly intermediates, potentially States G/H, accumulate in the nucleolus.

A fourth cluster (“Other”) comprised factors with no known direct role in ribosome assembly, including members of the NMD pathway, the exon junction complex, and the SF3b spliceosome complex. Phenotypically, this group showed reduced NPM1 dynamics and increased nucleolar intensity (Fig. 3D, S9A; further characterization in Supplementary Text and Fig. S9A-F). Despite their non-ribosomal functions, FISH revealed that depletion of SF3b factor PHF5A caused nucleolar accumulation of all three pre-rRNA intermediates, and Northern blot showed an increase in the 47S/45S/41S species, consistent with an early assembly stall (Fig. 4H, S6B, S9E). A similar but weaker effect was seen upon depletion of CNOT1, a core component of the CCR4-NOT deadenylase complex (Fig. S9B), suggesting that disruption of mRNA processing pathways converges unexpectedly on early ribosome biogenesis defects and accumulation of aberrant, early precursors in the nucleolus. Together, these results establish that distinct nucleolar biophysical states defined by coherent changes in NPM1 dynamics and nucleolar morphology correspond to the accumulation of specific pre-ribosomal intermediates at defined stages of ribosome biogenesis.

### NPM1 dynamics and nucleolar morphology reflect pre-ribosome intermediate composition

Across phenotypic clusters, changes in NPM1 dynamics were mirrored by changes in nucleolar morphology, where conditions associated with slower dynamics produced rounder, more intense, more cohesive condensates, while conditions associated with faster dynamics produced larger, irregular condensates with disrupted integrity. This coherent covariation suggests that both readouts report on a shared physical property of the nucleolus: the strength of interactions that hold the condensate together.

Current models in the field suggest that nucleolar condensation is driven by *trans* interactions between NPM1 and surfaces on ribosome intermediates, in particular rRNA and IDRs. The valency of *trans* interactions is thought to decrease as the ribosome matures, ultimately supporting thermodynamic flux of the ribosome out of the nucleolus [24, 30]. We therefore hypothesized that the strength of interactions between NPM1 and developing pre-ribosomes determines condensate cohesion and scaffold dynamics. In support of this idea, we observed a strong direct correlation between NPM1 partitioning and t_1/2_ for representative hits across fast and slow phenotypic clusters (Fig. 5A, r = 0.87, p = 0.0001). This trend suggests that conditions that drive stronger partitioning into the condensate also slow NPM1 exchange, while conditions that weaken partitioning accelerate it. Furthermore, increased irregularity (as measured by decreased circularity and increased eccentricity) correlated with shorter t_1/2_ across clusters (Fig. 3D), linking disruption of condensate integrity to faster molecular exchange. Together, these observations support a hypothesis that the valency of NPM1-pre-ribosome interactions sets the material state of the nucleolus by determining simultaneously how cohesive the condensate appears morphologically and how dynamically its scaffold exchanges.

**Figure 5:**
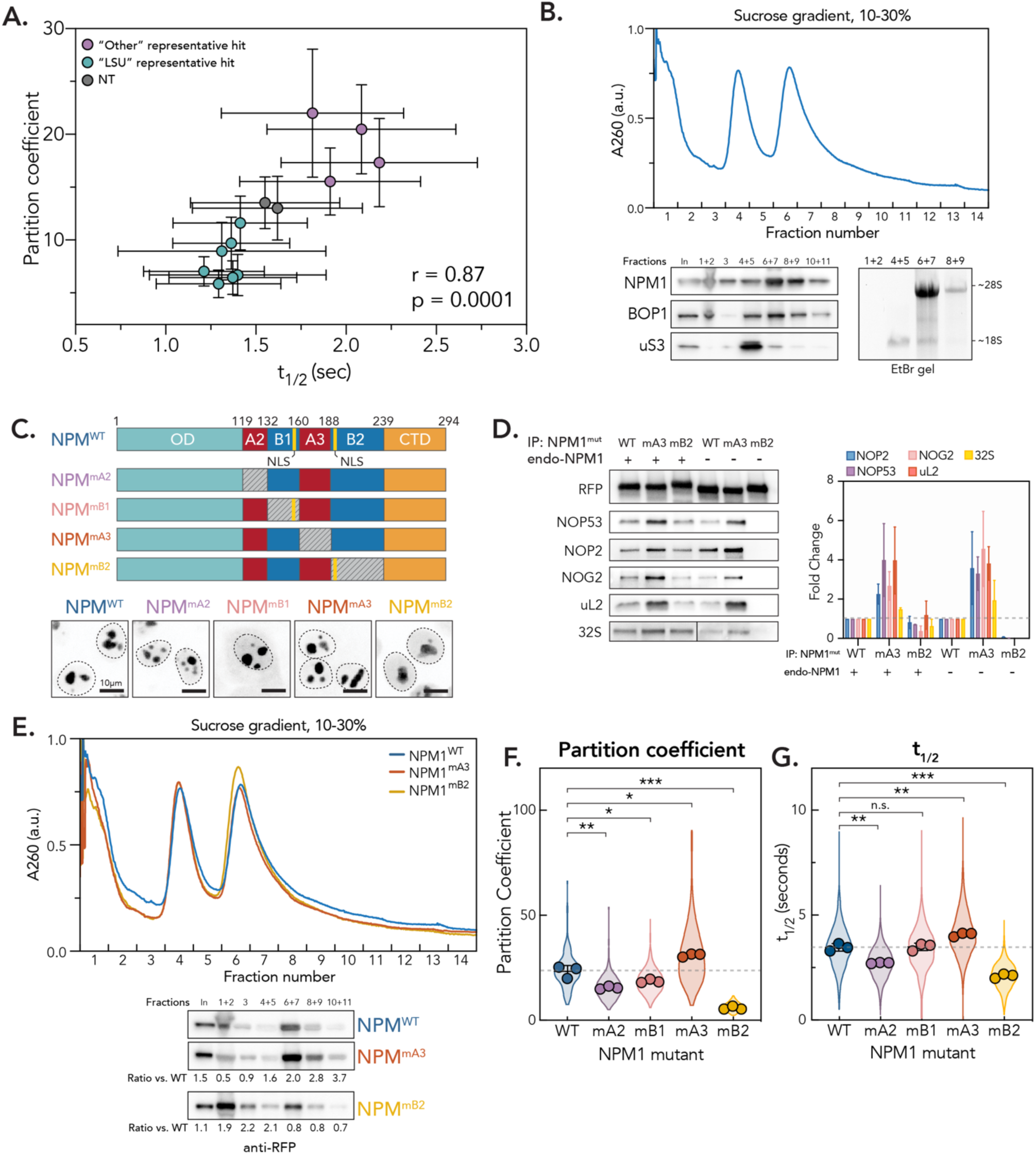
NPM1 dynamics are determined by interactions that drive partitioning into the nucleolus. (A) Partition coefficients vs. t_1/2_ for representative hits in phenotypic clusters (n = 100 nucleoli, points are mean, error bars are SD). Purple circles = representative Other” hits. Teal circles = representative “LSU” hits. Pearson r and p-value indicated. (B) Sucrose gradient fractionation of pre-ribosomal complexes from nuclear fraction of HeLa cells transduced with NPM1^WT^-mScarlet. Bottom left, western blot for indicated proteins shown below for pooled fractions. Ethidium bromide-stained RNA gel for total RNA isolated from indicated pooled fractions shown to bottom right. Approximate size of 28S and 18S indicated. (C) Schematic of NPM1 mutant constructs. Each construct is also N-terminally fused with mScarlet. Gray box represents charge neutralization. Yellow lines represent annotated nuclear localization signal (NLS). Representative images shown for localization pattern of each mutant construct (mScarlet signal). Dotted line outlines nucleus. Scale bar is 10 μm. (D) Left: representative western blot for ribosome assembly factors and proteins associated with immunoprecipitated NPM1 with indicated mutations. Northern blot for ITS2 showing 32S band shown at bottom. Right: Quantification of blots from IPs across replicates. Error bars are SD. n = 3. (E) Sucrose gradient fractionation of nuclear pre-ribosomal complexes isolated from HeLa cells transduced with NPM1^WT^-mScarlet (blue curve), NPM1^mA3^-mScarlet (orange curve), and NPM1^mB2^-mScarlet (yellow curve). Below, western blot showing mScarlet-tagged (RFP antibody) NPM1 from pooled fractions as indicated. Quantified ratio of RFP signal normalized to WT shown below each mutant blot. (F) Partition coefficients for each mutant construct. Violin plot shows spread of individual points for 3 biological replicates (n = 40 nucleoli per replicate, points show means of each replicate, error bars are SD). p-values calculated using two-tailed unpaired t-test between biological replicates. * p < 0.05, ** p<0.01, *** p < 0.001. (G) t_1/2_ for each mutant construct. Violin plot shows spread of individual points for 3 biological replicates (n = 250 nucleoli per replicate, points show means of each replicate, error bars are SD). p-values calculated using two-tailed unpaired t-test between biological replicates. n.s. = not significant, ** p<0.01, *** p < 0.001.

### NPM1 dynamics and nucleolar partitioning are determined by interactions with developing pre-ribosomes

To directly test whether NPM1-preribosome interactions dictate dynamics, we performed sucrose gradient sedimentation and found that NPM1 preferentially co-sediments with pre-LSU complexes (Fig. 4G and 5B), consistent with prior reports [66]. To ask how assembly state affects this association, we examined NOP53-depleted cells, where late LSU intermediates accumulate and NPM1 dynamics are accelerated (Fig. 4F-H). We found that NPM1 cosedimentation with the pre-LSU peak is markedly reduced in this context (∼75% decrease as compared to control, Fig. 4H), consistent with the prediction that late LSU intermediates, which have undergone partial rRNA folding and factor displacement, present fewer NPM1-binding surfaces. This result links the assembly state of pre-ribosomal particles directly to NPM1 association strength, supporting the idea that ribosome maturation progressively reduces the valency of condensate-scaffold interactions.

We then sought to identify the molecular features of NPM1 that mediate this maturation-sensitive association and thereby determine scaffold dynamics. NPM1 contains an N-terminal pentamerization domain, an intrinsically disordered linker region, and a C-terminal RNA-binding three-helix bundle (Fig. 5C). The IDR exhibits a charged block pattern, where two acidic domains (A2 and A3) that mediate heterotypic interactions with positively charged nucleolar IDRs alternate with two basic regions (B1 and B2) that bind to rRNA [23]. Homotypic interactions between A3 and B2 can also drive phase separation [22], although these appear to contribute less strongly to partitioning than heterotypic interactions [24]. To determine how these domains contribute to the interactions between NPM1 and pre-LSU complexes, phase separation, and dynamics in cells, we generated lentiviral constructs with mScarlet-tagged NPM1 where we charge neutralized each IDR region (NPM1^mA2^, NPM1^mB1^, NPM1^mA3^, and NPM1^mB2^, see Fig. 5C and Methods) and introduced them into unedited HeLa cells. To eliminate the contribution of heteropentamer formation with endogenous WT NPM1, we also introduced constructs into HeLa cells where endogenous NPM1 was stably ablated (Fig. S10A-C).

We assessed mutant interactions with pre-LSU complexes by co-IP western blot, qPCR, and ITS2 Northern blot, as well as sucrose gradient co-sedimentation. Mutation of A3 increased association with markers of pre-LSU intermediates and the 32S pre-rRNA as well as co-sedimentation with pre-LSU complexes, while loss of B2 led to decreased association (Fig. 5D and E and S10D-F). These impacts were strengthened in the absence of WT NPM1 (Fig. 5D and S10D-F). The increased preribosome association upon A3 removal supports in vitro findings that intramolecular A3/B2 interactions tune the availability of B2 for heterotypic rRNA binding, potentially acting as a regulatory switch^23^, and demonstrates that B2 is the primary driver of preribosome association.

Having established that IDR mutations directly tune NPM1-preribosome interactions, we asked how this impacts partitioning and dynamics. In the presence of endogenous NPM1, NPM1^mA2^ and NPM^1mB1^ showed modest decreases in nucleolar partitioning (∼30% and ∼20%, respectively), consistent with loss of heterotypic interactions with R-rich proteins and rRNA. Neutralization of A3, however, increased partitioning by ∼30%, while loss of B2 caused a ∼75% decrease, with NPM1^mB2^ largely excluded from the nucleolus in the knockout background (Fig. 5F, S10C). Strikingly, these changes in partitioning were directly reflected in NPM1 dynamics (Fig. 5G). The strongly partitioning construct NPM1^mA3^ showed a ∼20% increase in t_1/2_, while NPM1^mA2^ and NPM1^mB2^, which partitioned more weakly, exhibited significantly faster recoveries (∼20% and 40%, respectively). These results directly correlate with NPM1-preribosome interaction strength (Fig. 5E). Together, these results demonstrate that the strength of NPM1-preribosome interactions determines both nucleolar partitioning and scaffold exchange dynamics (Fig. 6A), providing a direct mechanistic basis for the changes in dynamics and morphology we observe across phenotypic clusters in our screen.

**Figure 6:**
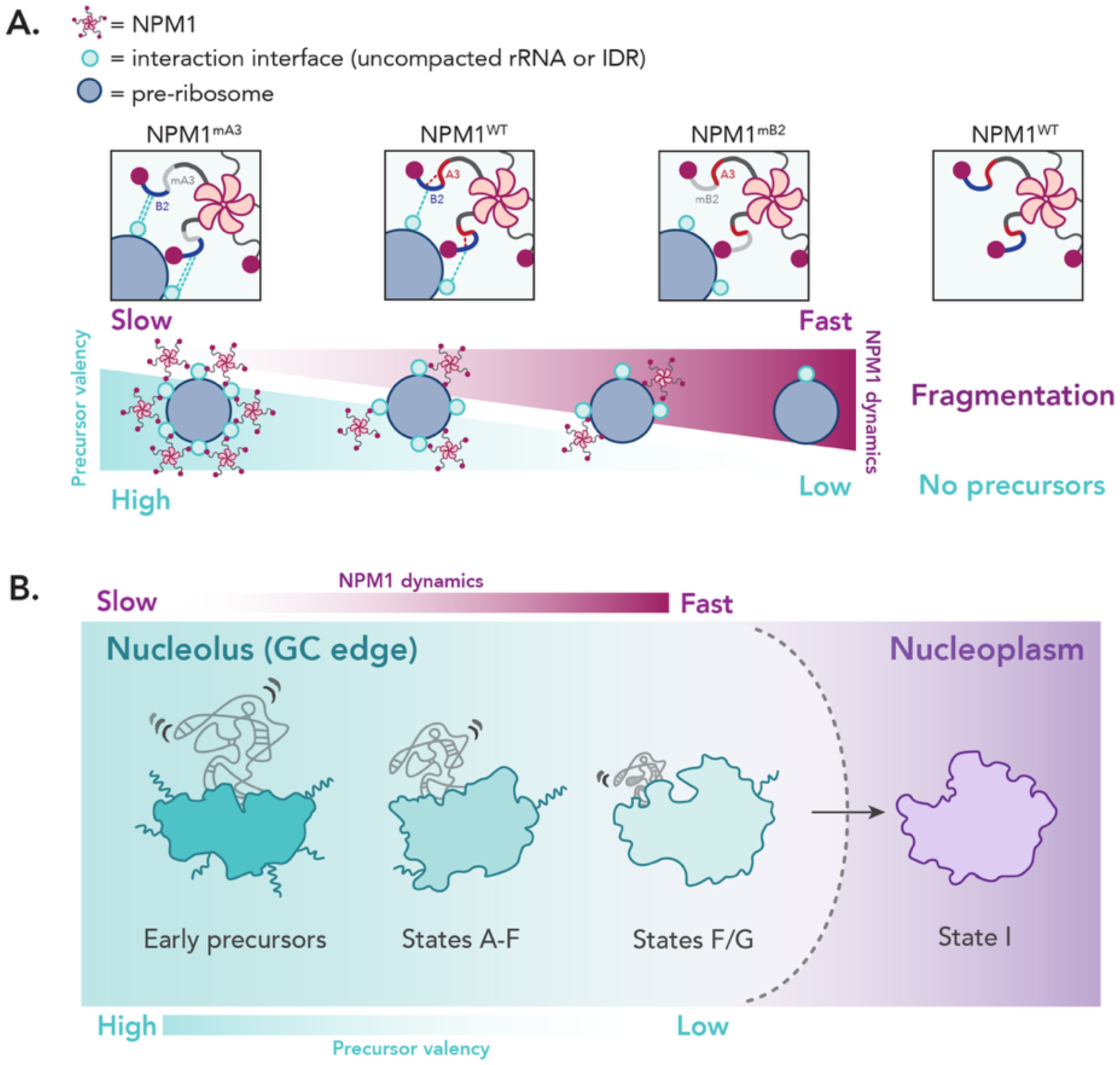
Model for coupling ribosome assembly to nucleolar dynamics. (A) Model for how ribosome intermediates determine nucleolar biophysical properties. Insets show how regions of the NPM1 IDR influence interactions with ribosomal precursors. Heterotypic interactions with surfaces on developing pre-ribosomes (represented by teal ball) shown as teal dotted line, homotypic interactions between A3 and B2 shown as red dotted line. (B) Model for pre-ribosomal maturation at the GC interface as it relates to nucleolar dynamics and precursor valency. Uncompacted IDRs (teal lines) and rRNA (gray lines) indicated graphically, with progressive loss over maturation.

## Discussion

Here, we use HiT-FRAP to show that NPM1 dynamics and nucleolar morphology are jointly sensitive to the assembly state of ribosomal precursors, and that this sensitivity reflects the valency of NPM1-preribosome interactions. Conditions that drive accumulation of high-valency early intermediates simultaneously slow NPM1 exchange and compact the condensate, while accumulation of low-valency late LSU intermediates accelerates exchange and disrupts condensate integrity. Importantly, mutations in the NPM1 IDR that strengthen pre-ribosome binding slow exchange and increase partitioning, while those that weaken binding accelerate exchange and reduce partitioning, providing direct mechanistic support for this model. Together, these findings demonstrate that the interaction networks driving nucleolar phase separation also determine the dynamic and morphological properties of the condensate, connecting ribosome assembly state to nucleolar material properties at the molecular level.

### Insights into nucleolar condensate biophysics and assembly principles

Our results support a physical picture where nucleolar scaffold dynamics and morphology are determined by the valency and thermodynamic strength of heterotypic interactions that drive condensate formation, as seen in other network fluids and in vitro reconstituted condensate systems [7, 67]. Therefore, dynamics and morphology together report on material state of the condensate, which is coupled to pre-ribosome assembly state. Importantly, this finding also indicates that the nucleolus does not behave as a simple fluid, which was historically inferred from studies using NPM1 as a proxy for liquid-like nucleolar behavior. This highlights a key question: how do individual nucleolar components contribute to the physical features of the condensate, and how do those features in turn shape condensate function?

In support of this question, it was recently shown that nucleolar proteins are substantially more dynamic than the rRNA gel they surround [34], consistent with the idea that a kinetically accessible environment facilitates chaperone-mediated rRNA folding and the sequential assembly factor exchanges required for modular maturation. Our results suggest that this kinetic accessibility may be set, at least in part, by the valency networks driving phase separation. If these networks determine NPM1 exchange rates, we speculate that they may similarly regulate the accessibility of the condensate environment to assembly factors more broadly. Systematically mapping how the dynamics of other nucleolar constituents relate to their function and to overall condensate material properties will be an important future direction, which we anticipate HiT-FRAP will enable.

### Connecting nucleolar assembly principles to ribosome assembly

Interestingly, it has recently been demonstrated that the nucleolus exhibits non-uniform material properties radially, with a more solid-like core and more liquid-like periphery [68]. Our results suggest a molecular basis for this gradient, where the valency of interactions between nucleolar scaffolds such as NPM1 and developing ribosomes dictate condensate material properties: less assembled precursors participate in more interactions with NPM1 to create more solid-like features, while more assembled intermediates are more compact and thus have less interfaces available for binding, resulting in increased fluidity. We hypothesize that the nucleolar surface most closely represents the liquid-like system represented in current NPM1-based nucleolar reconstitution systems. Therefore, we propose that NPM1 dynamics may most strongly impact late subunit maturation and nucleolar export at this interface, which we discuss further below.

Consistent with previous results [35, 38, 66, 69], we find that disruption of late LSU assembly leads to nucleolar accumulation of late abortive intermediates, likely human States G/H, which weakens the nucleolar condensate and leads to disrupted morphology and increased NPM1 exchange. We therefore propose that the human State G/H intermediates represent the final stages before export, where the transition to State I licenses release by falling below the threshold of valency required for nucleolar sequestration. This transition is hallmarked by installation of the CP coupled with binding of the rixosome, a complex necessary for ITS2 processing. *Trans* interactions may change in several ways during this transition. Several IDRs are lost or stabilized, although in total, IDRs increase due to arrival of rixosome components (Fig. S11). More notably, domains IV and V of the 28S rRNA mature, collectively resulting in a ∼30% decrease in uncompacted rRNA (Fig. S11). This mechanism appears to be conserved, as recent work in yeast identified a similar LSU intermediate as the final nucleolar assembly stage [29]. However, intriguingly, engineered rRNA that lacked large portions of ITS2 and 28S expansion segments did not globally impact ribosome export, despite being comparable in size to the compaction of the 28S that occurs during the transition to State I [16, 70, 71]. These findings suggest at least two possibilities: 1) that certain “uncompacted” rRNA regions may not be available for *trans* interactions, or 2) that additional factors outside of these interactions may govern sequestration within the nucleolus and export to the nucleoplasm. Therefore, further work characterizing export incompetent intermediates will be necessary to understand the structural mechanism for nucleolar release.

Taken together, we propose that the biophysical features and ribosomal intermediates associated with the “LSU” group represent the outermost molecular “layer” of the nucleolus, providing biophysical insight into the nucleolar interface (Fig. 6B). This idea is supported by recent *in situ* structural efforts in *Chlamydomonas* nucleoli that revealed a gradient of ribosome assembly intermediates at the nucleolar surface [72]. Importantly, interfaces are a unique environment where the condensate and its surroundings meet, and as such, have been shown to form distinct chemical and functional environments in other condensate systems [73–75]. In the case of the nucleolus, the final stages of precursor assembly must interact with the nuclear environment, which could influence these steps in nontrivial ways. Consistent with this idea, it has been shown that disruption of chromatin structure leads to amorphous nucleolar morphology reminiscent of the LSU phenotype [34, 76]. It is therefore likely that the nucleolar interface possesses specialized features to facilitate handoff. In support of this hypothesis, recent work has shown that the nucleolar rim is a compositionally distinct compartment, and intriguingly, is enriched for NPM1, factors removed in the State F to G transition (BOP1, DDX18, EBP2, FTSJ3, NOC2L, NOC3L), State G/H factors (NOG2, RPF2, uL18), and factors involved in remodeling of State G/H to I (GNL3, CCDC86) [77]. It will be exciting to consider how the unique environment of the nucleolar interface may facilitate the key molecular events associated with these architectural changes, while remaining distinct from the nuclear surroundings.

Lastly, given that we observe such striking disruption of nucleolar biophysical features upon accumulation of abortive intermediates, we speculate that nucleolar assembly principles could be linked to ribosomal quality control. We believe our results support the thermodynamic gating model previously proposed in the literature [24, 30, 63], where only properly assembled intermediates that have undergone the compaction and remodeling necessary to fall below a critical valency threshold are competent for export, while aberrant or stalled particles remain sequestered. Importantly, however, our results suggest that the dynamic, liquid-like character of the nucleolar surface may be essential for this gate to function, by ensuring that export-competent particles can traverse the interface on biologically relevant timescales rather than remaining kinetically trapped, while incompetent particles remain in the interior.

### Biophysical changes in the nucleolus in response to stress

We also discovered unexpected ties between nucleolar dynamics and diverse mRNA processing pathways. Connecting our observations to reports in the literature suggest that they may relate to the nucleolus’s role in cellular stress responses. For example, rRNA processing stalls during acute stress and leads to accumulation of unprocessed precursors, which decrease nucleolar dynamics and are stored until stress is alleviated [44, 78]. This response is reminiscent of the pre-mRNA processing phenotype we observe and suggests it may result from induction of cellular stress. In support of this idea, disruption of splicing causes stress granule formation and proteotoxic stress^81,82^. Furthermore, during stress, the nucleolus reversibly stores misfolded proteins whose aggregation is prevented through interactions with scaffolds such as NPM1 [43]. This accumulation leads to nucleolar rigidification, similar to the “Other” phenotype. Interestingly, inhibition of NMD induces expression of truncated proteins that aggregate and could enter the nucleolus [79], potentially leading to the rigidification we observe. Therefore, the “Other” phenotype may be a biophysical signature of nucleolar stress. If so, nucleolar dynamics may serve as a sensitive integrator of cellular homeostasis, with implications for diseases in which nucleolar function and RNA metabolism are co-disrupted, including ALS and FTD, where NPM1 dynamics, nucleolar stress, and RNA processing pathways are all perturbed [32, 80–82].

## Conclusion

Our work establishes that the assembly state of ribosomal precursors determines the material properties of the nucleolar condensate through the valency of interactions with the scaffold NPM1, connecting ribosome biogenesis to nucleolar biophysics at the molecular level. This model implies that the same interaction networks that drive phase separation also govern condensate dynamics, suggesting that material properties may be directly tunable by the composition and connectivity of condensate interaction networks [83]. This principle may be relevant to pathological condensates, where shifts in the stoichiometry of key interactors could drive transitions toward poorly dynamic, aggregated states [1]. More immediately, our findings raise the question of how NPM1 dynamics influence late LSU maturation and nucleolar export, which has direct relevance to diseases that disrupt nucleolar interaction landscapes. More broadly, our results raise the possibility that condensate material properties are not merely downstream consequences of the interactions they house but are themselves part of the functional machinery, actively shaping the kinetic landscape of the processes they organize. HiT-FRAP now provides a platform for testing this principle systematically, both in the nucleolus and across other biomolecular condensates whose dynamic properties remain mechanistically uncharacterized.

## Author contributions

Conceptualization, J.S-G.; Investigation, J.S-G., R.O.O., S.L.; Writing – Original Draft, J.S-G.; Writing – Review & Editing, J.S-G., S.N.F., R.D.V.; Methodology, J.S-G., X.Y., N.S.; Resources, J.S-G., S.N.F., and R.D.V.; Formal analysis, J.S-G., X.Y., N.S.; Software, J.S-G., X.Y., N.S.; Funding Acquisition, J.S-G., S.N.F., and R.D.V.; Supervision, J.S-G., S.N.F. and R.D.V.

## Acknowledgments

We thank members of the Floor, Vale, and Sheu-Gruttadauria labs for feedback on this work and their continued support. We additionally thank Dyche Mullins and members of the Mullins lab for their generous support. SMG1i was generously provided by Robert Bridges, Ph.D. of the Rosalind Franklin University of Medicine and Science and Cystic Fibrosis Foundation Chemical Compound Program. JS-G is a Hanna H. Gray Fellow of the Howard Hughes Medical Institute. This work was supported by the National Institutes of Health R35GM149255 (to SNF). SNF is a Pew Scholar in the Biomedical Sciences, supported by The Pew Charitable Trusts. RDV is supported by the Howard Hughes Medical Institute.

## Competing interests

None declared.

## Materials and Methods

## Key resources table

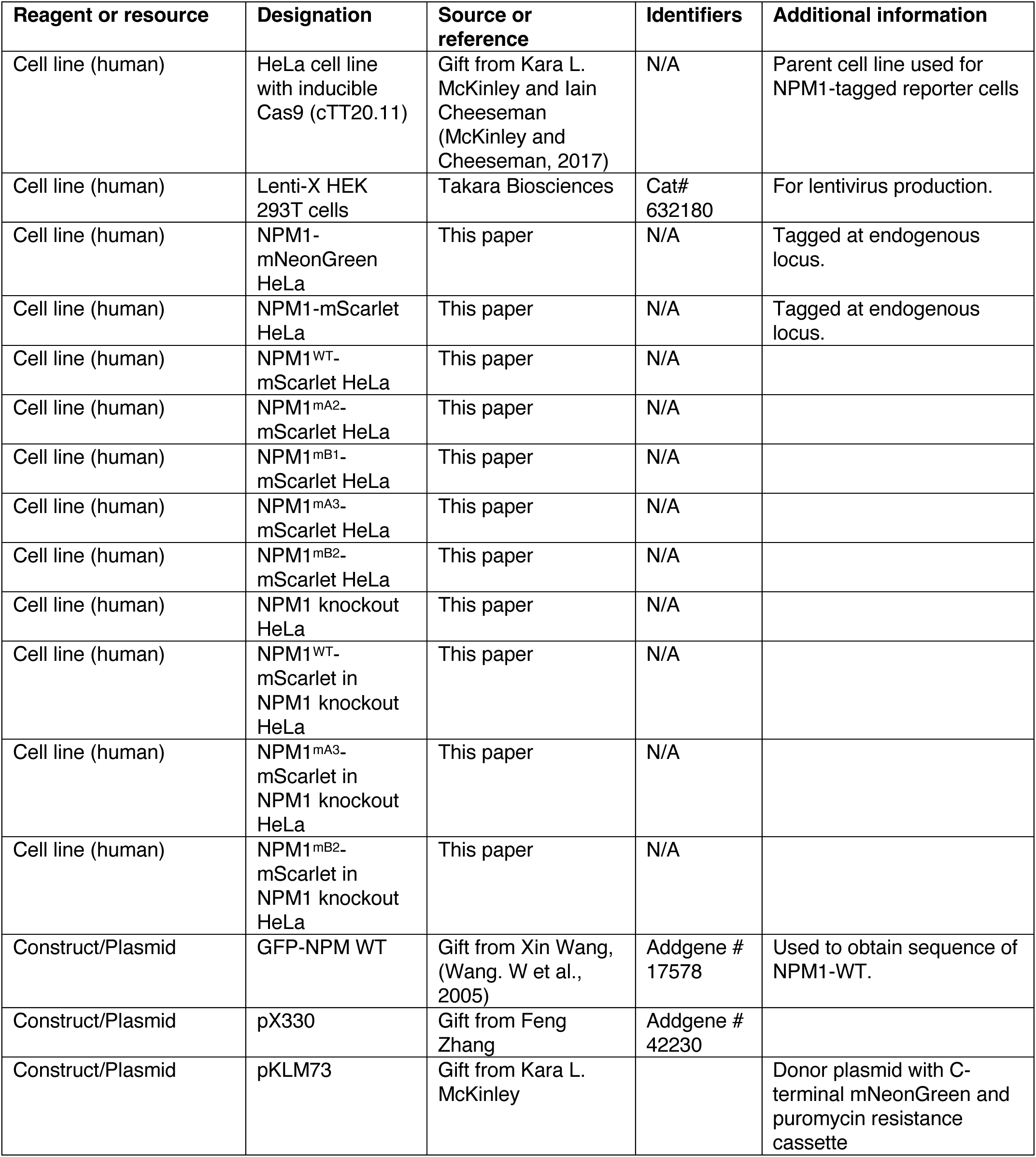
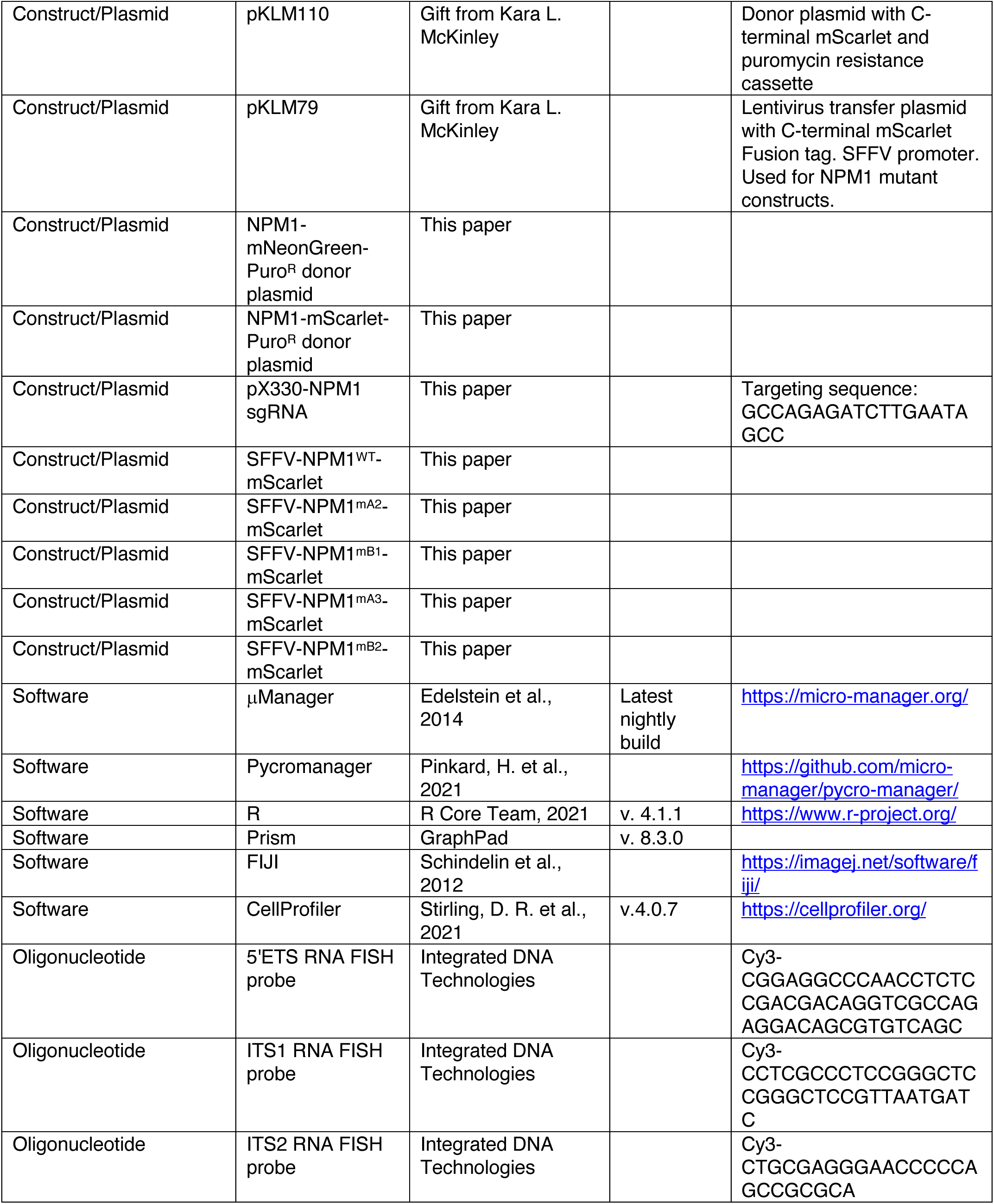
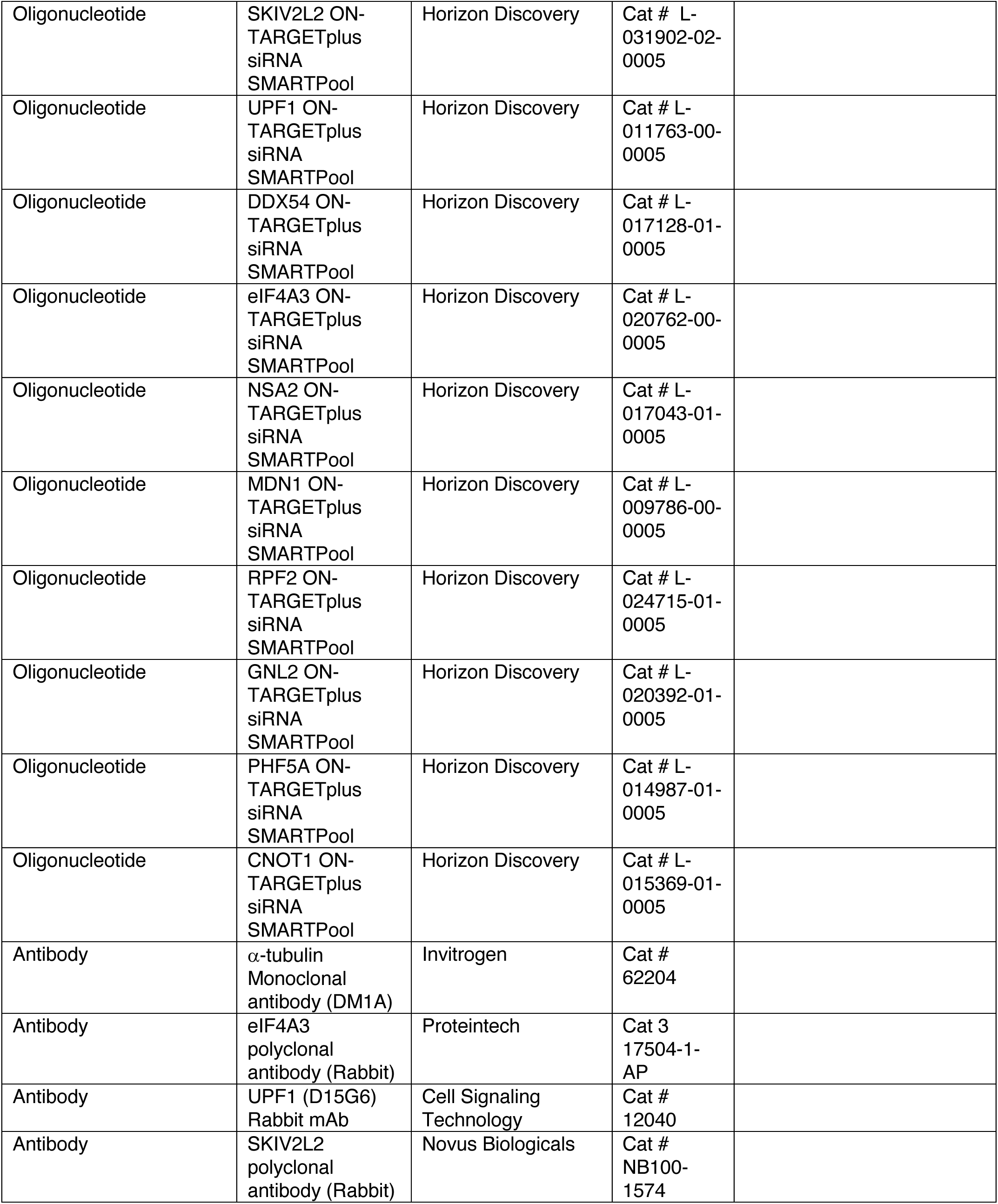
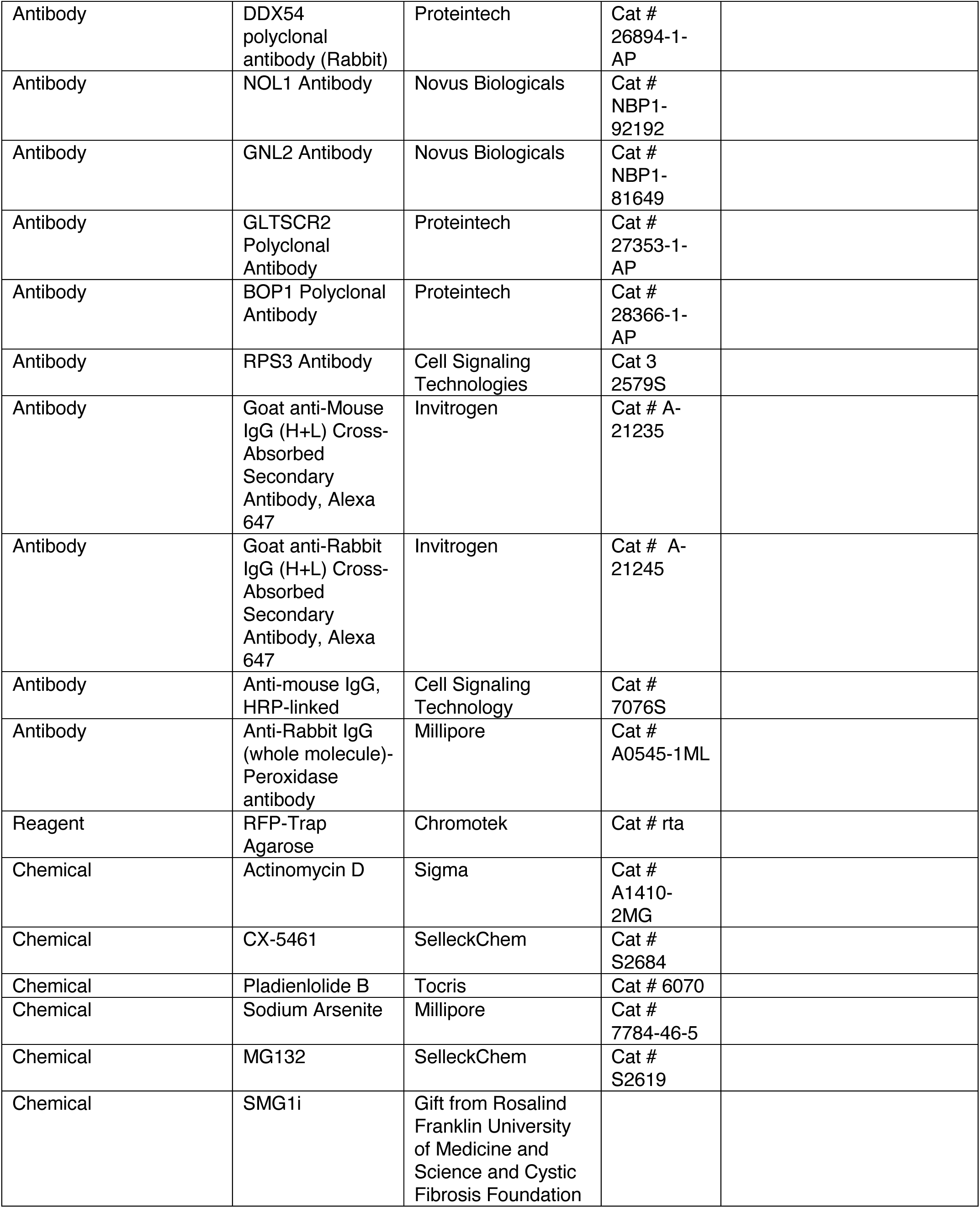
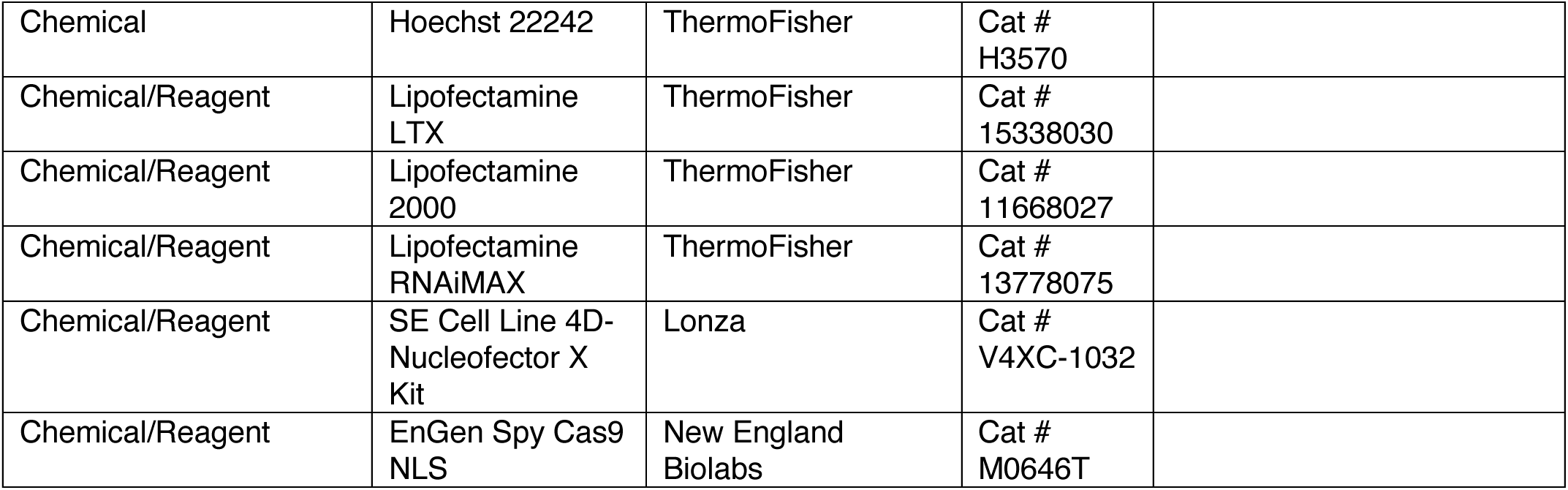

### Cell culture

HeLa (cTT20.11 [84], gift from Kara L. McKinley and Iain Cheeseman) and Lenti-X HEK293T cells (Takara Bio, used for lentivirus production) were cultured in DMEM (Gibco) supplemented with 10% tetracycline-free FBS (Gemini) and penicillin and streptomycin (Gibco) at 37°C with 5% CO_2_ in a humidified incubator. Cells were maintained at passage number under 25 and routinely checked for mycoplasma. Cells were passaged using 0.05% Trypsin-EDTA (Gibco).

### Endogenously-tagged cell line generation

Endogenously-tagged cell lines were generated using CRISPR-Cas9 homology directed repair. Guide RNAs targeting the last exon of NPM1 and FBL were designed using the Benchling CRISPR tool to maximize on-target and minimize off-target effects (targeting sequence: NPM1-GCCAGAGATCTTGAATAGCC; FBL-AACTGAAGTTCAGCGCTGTC). This sequence was inserted into pX330 (gift of Feng Zheng, Addgene #42230), which encodes both spCas9 and the sgRNA. Homology arms for genes of interest were ordered from IDT as gBlocks with PAM sequence for sgRNA mutated. Homology arms were cloned into pKLM73 for mNeonGreen and pKLM110 for mScarlet (gifts from Kara L. McKinley) using NEBuilder HiFi DNA Assembly (New England Biolabs). To generate knock-in lines, HeLa cells were seeded in a 6-well plate at 1×10^6^ cells per well. The following day, cells were transfected with 1 µg each of the pX330 and donor plasmids using Lipofectamine 2000 (ThermoFisher). After 72 hr, cells were passaged into 10 cm dishes and cells with integration were selected for using puromycin at 1 µg/mL. After sufficient cell death and outgrowth of resistant colonies, cells were passaged one additional time to recover and then fluorescently positive cells were sorted for single cells into a 96-well plate using a Sony SH800 cell sorter. Single-cell clones were checked for correct localization of tagged protein and confirmed by genotyping PCR. Briefly, gDNA was isolated from a confluence 12-well plate by rinsing once with PBS followed by addition of 0.5 mL genomic DNA lysis buffer (100 mM Tris pH 8, 5 mM EDTA, 200 mM NaCl, 0.2% SDS, 0.2 mg/mL proteinase K). Plates were incubated with lysis buffer at 37°C overnight. Lysed cells were transferred to an Eppendorf tube and DNA was precipitated using isopropanol followed by washing with 70% ethanol. Pellet was resuspended in TE buffer (1 mM EDTA pH 8, 10 mM Tris pH 8) and used as a template for amplification of insertion regions. Products were visualized for size differences by agarose gel and then gel purified and submitted for Sanger sequencing.

### Knockout cell line generation

NPM1 was stably ablated from HeLa cells (cTT20.11) using CRISPR-Cas9 as follows. Gene Knockout Kit v2 against NPM1 from Synthego was used for guide RNAs. 180 pmol guides were complexed with 20 pmol EnGen Spy Cas9 NLS (New England Biolabs) for 10 min at room temperature. The assembled Cas9 RNP was nucleofected into 1.5 x 10^5^ cells using Lonza 4D-Nucleofector X unit and SE Cell Line Kit following manufacterer’s protocol. Polyclonal population was assessed for knockdown by western blot and then sorted into single cells using a SONY SH800. Single cell clones were assessed for knockdown by western blot against NPM1 and Sanger sequencing.

### Microscopy

All live-cell and fixed-cell confocal imaging was carried out on a Nikon Ti-E inverted microscope equipped with a Yokogawa CSU-X high-speed confocal scanner unit and a pair of Andor iXon 512 × 512 EMCCD cameras. All components of the microscope were controlled by the open-source platform µManager [85]. The microscope stage was enclosed in a custom-built incubator that maintains preset temperature. Live-cell imaging was performed at 37°C with 5% CO_2_ and cells were equilibrated on the microscope for at least 30 min prior to imaging. Fixed-cell imaging was performed at room temperature. High-magnification images were obtained using a 100x 1.49 NA oil-immersion objective. HiT-FRAP and high-content images were acquired with a 40x 0.95 NA air objective. For HiT-FRAP, images for mNeonGreen cells and mScarlet cells were collected at 100x EM gain with a 50 ms and 100 ms exposure, respectively. Bleaching was performed with a 405 nm focused laser beam set at 10 mW power steered by a pair of galvo mirrors (Rapp UGA-40), which was controlled by the Projector Plugin in µManager.

### HiT-FRAP acquisition

Automated FRAP experiments were performed using custom code and the Python library Pycromanager [86], which integrates hardware control with image processing via µManager. Image processing was performed using scikit-image [87]. Positions and wells for acquisition were defined using the µManager High Content Screening (HCS) Site Generator Plugin. Exposure length was predetermined for each cell line to avoid saturated pixels. HiT-FRAP acquisition proceeded as follows: at each field of view, a single image was acquired. Organelles of interest were then identified. Firstly, the background pixel threshold was determined using Otsu’s method. Threshold values for nucleoli were determined using local thresholding, and regions larger than 1000 pixels were removed. Both thresholds were combined and then subjected to two rounds of erosion and dilation, followed by removal of objects smaller than 10 pixels to clear the background. To maximize throughput, further analysis and acquisition only continued if greater than 20 organelles were present in a field of view. Centroids for organelles were then determined to generate a bleach position list. We set a size threshold for bleached organelles of at least twice the area of the bleach spot to preferentially capture internal mixing rather than exchange with the nucleoplasm, although due to limitations in magnification and the fixed size of our bleach laser, whole-organelle bleaches could not be completely avoided. A random 50% of these positions were then chosen for bleaching, with the remainder kept as controls to account for acquisition photobleaching. Bleach locations were exported for every field of view to be used in post-analysis. Acquisition then started, followed by concurrent bleaching at designated organelle centroid positions. Length of acquisition was manually pre-determined to achieve recovery plateau. This process was iterated through multiple fields of view until 500 bleaches were acquired, after which the stage was moved to the next well and the sequence repeated.

To account for drift in the bleach laser position, automated recalibration was performed as follows: after every two fields of view, a single image was acquired. The image was thresholded to identify cells using the triangle algorithm, and a position closest to the center that did not have a foreground object (cell) was determined. The bleach laser was then targeted to this location and an image was taken of the exposure. The true bleach laser position was determined from this image by identifying the maximum intensity position in the image. To increase the speed of finding this location, a region of interest was used based upon the estimated location of the bleach laser. When the square of the distance between the intended target and the true bleach spot location was greater than an offset of 5, a full calibration of the laser was performed.

### HiT-FRAP image analysis

Analysis of high-content FRAP data was performed using the same software used for acquisition. For each FRAP recovery time series, nucleoli were identified in frame 0 using the same thresholding procedure as used for acquisition. Bleach spots were then identified as follows: the bleach positions generated during acquisition (pointers) were linked to the bleach frame. To account for experimental offsets in bleach positions, true bleach locations were determined empirically. Briefly, it was manually determined that the minimum intensity post-reported bleach time occurred after four frames (bleach time offset). For each pointer position, the minimum intensity frame (bleach frame + bleach time offset) was subtracted from the bleach frame and smoothed to remove background noise. Otsu global thresholding was then applied to identify bright spots (detected bleach spots). After two rounds of erosion and dilation to minimize background noise, the centroids of the closest detected bleach spots to each original pointer position were used as the true bleach location. These true bleach coordinates were then linked to the corresponding organelle. Bleach spots that fall outside of a nucleolus, bleach the same nucleolus, overlap, or are too far away from the original pointer position (>20 pixels in any direction) were filtered. Bleach spot masks were then generated by subjecting the true bleach coordinates to three rounds of dilation.

Using these bleach spot masks, FRAP curves were determined as follows: to monitor photobleaching due to acquisition, control spot masks were generated by identifying centroids of unbleached organelles and submitting these to three rounds of dilation. Raw intensities for the bleach and control spots were calculated over the time course. Background was determined for each frame by generating a binary image of regions with a pixel intensity of less than 300, removing regions smaller than 50 pixels. The mean intensity of the largest area was used as background and subtracted from all measured intensities. A photobleaching factor was then determined for control spots and calculated as the intensity ratio between the control spot at every time point as compared to intensity at t = 0. The mean of this ratio across all control spots was then used as a correction factor for bleach spot intensities. These intensities were then normalized to the pre-bleach intensity and minimium intensity (0 to 1) to generate final intensities for fitting. These intensities were then fit to several FRAP models (single exponential, double exponential, soumpasis, and ellenberg), although single exponential fittings were used in all subsequent analyses. Curves were filtered out if they had less than 5 frames prior to bleach (to avoid poor determination of pre-bleach intensity), imaged for less than 100 frames post bleach for mNeonGreen or 50 frames for mScarlet (to ensure adequate recovery), if the mobile fraction was less than 0 or greater than 1.05, and if R^2^ < 0.7. Parameters from the fitted curve are then exported (initial slope, mobile fraction, and t_1/2_).

Organelle features were determined as follows: for each bleached nucleolus, the following organelle features available in scikit-image were calculated: area, mean intensity, circularity, eccentricity, solidity, minimum intensity, maximum intensity, perimeter, hu moments (2^nd^, 3^rd^, and 4^th^), intensity percentiles (1^st^, 5^th^, 95^th^, and 99^th^), median intensity, the standard deviation of intensities, intensity kurtosis, and intensity skew.

Analysis code for HiT-FRAP acquisition and analysis is publicly available on GitHub (https://github.com/jess-sheu/colony_blob_bleacher) and archived on Zenodo (https://doi.org/10.5281/zenodo.20275447).

### Dynamics screen phenotype analysis

Gene depletions with high cell death were removed manually from all subsequent analysis (DDX19B, DDX19A+DDX19B, DDX39A+DDX39B, and SNRNP200, CRNKL2, CWC22, EFTUD3, eS4, eS6, PRPF8, SNW1). Dynamics features (mobile fraction and t_1/2_) were normalized using the median and median absolute deviation for all non-targeting siRNA controls in the same plate to account for plate-to-plate variation (robust z-score). Medians of robust z-scores were computed and used in final hit calling. p-values were calculated using the Kolmogorov-Smirnov test as compared to a randomly chosen subset (10%) of non-targeting control wells. FDR was determined using the Benjamini-Hochberg procedure. As indicated in the figure legends, an FDR threshold of 0.05 was used for defining significance. Thresholds for z-scores were set as indicated in figure legends.

### Principal component analysis for multidimensional analysis

Phenotype scores were calculated and incorporate the -ln(p-value) (determined by Kolmogorov-Smirnov test) and effect size as compared to a subset (randomly selected 10%) of non-targeting control wells in the same plate to account for plate-to-plate variation. The remainder of non-targeting controls were treated as experimental samples to test the robustness of the scoring method. Phenotype scores were scaled from 0 to 1 prior to further analysis. PCA was performed using the prcomp R package. The R packages ggplot2 [88], ggfortify [89], and ggrepe [90] were used to generate figures and are available through CRAN. Hits were manually called by separation from NT control cluster.

### Hierarchical clustering to identify phenotype clusters

Robust z-scores for hits were used in hierarchical clustering using the pheatmap R package [91] by Ward’s minimu variance method, variant D2. The resulting dendrogram was split into subtrees until clusters represented similar overall trends, generating five phenotypic clusters shown in Fig. S4B.

### Cloning for mutant NPM1 lentivirus constructs

All constructs were cloned into the lentiviral transfer plasmid pKLM79 (gift of Kara L. McKinley), which contains a C-terminal mScarlet fusion tag. The wild-type sequence of NPM1 was cloned from GFP-NPM WT (gift from Xin Wang, Addgene plasmid # 17578). Mutant NPM1 plasmids were constructed by ordering mutated regions as gBlocks from IDT with overhangs complementary to proximal regions. Proximal regions were then amplified from the wild-type construct to generate fragments with appropriate overhangs for assembly by NEBuilder HiFi DNA Assembly (New England Biolabs) into pKLM79 (cut with MluI and SpeI, New England Biolabs).

### Lentivirus transduction

Lentiviral particles were packaged by transfecting Lenti-X 293T cells with transfer construct of interest and PSP and VSVG helper plasmids using Lipofectamine LTX (Thermo Fisher) reagent. Media was changed 18 hr later and virus was harvested 72hrs after transfection. Virus was cleared by passing through a 0.45 µm PVDF filter. 200 µL viral supernatant was used to transduce a 6-well plate of HeLa cells by spinfection. Briefly, cells were seeded in suspension at 250,000 cells per well in the presence of viral supernatant and polybrene (Sigma-Aldrich) at a final concentration of 10 µg/mL. Cells were spun at 1000 x g for 45 min at 37°C and allowed to recover overnight. Media was replaced the following day and transduction was allowed to proceed for 48 hr until passage. Cells were passaged for at least two passages to eliminate cells that exhibited toxic levels of expression. Fluorescent cells were sorted using a Sony SH800 cell sorter gated to select the top 40% of expressing cells. Polyclonal populations were used for all experiments.

### siRNAs

For the primary RNA helicase and secondary screen, candidate libraries of ON-TARGETplus SMARTPool siRNAs (4 guides per gene) were ordered in arrayed format from Horizon Discovery. Non-targeting siRNA pools were interspersed randomly throughout the plate at a ratio of 1:6 to on-target pools. For depooled validation, deconvoluted guide pools (single guides) were ordered in an arrayed format for select hits. Single non-targeting guides were interspersed randomly throughout the plate at a ratio of 1:6. For knockdown of single genes, ON-TARGETplus siRNA pooled guides were ordered from Horizon Discovery (see Key Resources Table).

### siRNA transfections

For screens and depooled validation experiments, siRNAs were reverse transfected at a final concentration of 5nM using Lipofectamine RNAiMAX reagent (Life Technologies). Briefly, siRNA:lipid mixes were pipetted into wells of a 384-well glass bottom dishes (Matriplate; Brooks). After a 15 min incubation, cells were seeded at 900 cells per well. Knockdown was allowed to proceed for 72 hr before imaging. For validation experiments (RNA FISH, IF, and single gene knockdowns), pooled guide RNAs were reverse transfected at a final concentration of 25 nM into 96-well glass-bottom imaging plates (Matriplate; Brooks), with cells seeded at a density of 8000 cells per well. For qPCR and westerns, cells were reverse transfected with pooled guide RNAs at a final concentration of 25 nM into 24-well tissue culture dishes (Corning) and cells were plated at 24,000 cells per well.

### Drug treatments

For all drug treatments, cells were seeded at 50,000 cells per well into 96-well glass bottom imaging dishes (Matriplate; Brooks) 24 hr prior to experiments. For ATP depletion, cells were washed twice in DMEM without glucose (Gibco) and then incubated for 10 min with 10 mM sodium azide and 6mM 2-deoxyglucose diluted in DMEM without glucose supplemented with 10% tetracycline free FBS (Gemini). For sodium arsenite, cells were treated with 200 µm sodium arsenite diluted in culture media for 1 hr prior to imaging. For MG132, cells were treated with 10 µm MG132 diluted in culture media for 1 hr prior to imaging. For Pol I inhibitors, cells were treated with 0.04 µg/mL actinomycin D (Sigma) or 500 nM CX5461 (SelleckChem) diluted in culture media for 2 hr. For pladienolide B treatment, cells were treated with 10 nM PladB (Tocris, initially diluted in DMSO) diluted in culture media for 24 hr. For SMG1i treatment, cells were treated with 300 nM SMG1i (initially diluted in DMSO) diluted in culture media for 24 hr. All drug treatments were compared to cells treated with equal volumes of vehicle. For all imaging experiments, cells were transferred to heated and equilibrated microscope at least 30 min prior to imaging.

### RNA fluorescent in situ hybridization (RNA FISH)

RNA FISH was performed in a 96-well plate format (see siRNA transfections). After knockdown or drug treatment, cells were fixed in 4% Paraformaldehyde in PBS at 37°C for 10 min. Cells were then permeabilized by incubation with ice-cold methanol at -20°C for 10 min. Cells were incubated with 1 M Tris pH 8 for approximately 10 min. 5’ Cy5 labeled DNA probes (see Supplementary Table 1) were ordered from IDT and resuspended at 1 µg/mL in water. Stock probes were diluted 1:1000 in RNA FISH hybridization buffer preheated to 42°C (15% deionized formamide, 10% dextran sulfate, 2x SSC from Ambion, 0.02% BSA, 0.2 mg/mL Baker’s yeast tRNA from Ambion). Plate was placed in plastic bag with wet paper towels and incubated at 42°C for 1 hr. Cells were washed with 2x SSC three times at room temperature, followed by incubation with Hoechst 33342 DNA dye diluted at 1:1000 in 2x SSC for 10 min at room temperature. Cells were washed a final three times with 2x SSC and then imaged.

### Immunofluorescence (IF)

Immunofluorescence was performed in 96-well plate format (see siRNA transfections). After knockdown or drug treatment, cells were fixed in 4% Paraformaldehyde in PBS at 37°C for 10 min. Cells were then permeabilized by incubation with ice-cold methanol at -20°C for 10 min. Fixed cells were blocked in 3% BSA in PBST (PBS from Gibco supplemented with 0.1% Tween-20) for at least 1 hr, followed by incubation overnight with primary antibodies diluted at 1:500 in antibody dilution buffer (3% BSA, PBST, 0.02% sodium azide). The following day, cells were washed three times with PBST followed by incubation for 1-2 hr at room temperature with appropriate secondary antibody conjugated to Alexa 647 (ThermoFisher) diluted at 1:1000 in antibody dilution buffer. Cells were then washed three times with PBST and incubated with Hoechst 33342 DNA dye (Invitrogen) diluted at 1:1000 for 10 min at room temperature. Cells were washed a final three times with PBST and then imaged.

### NPM1 Immunoprecipitation for western blotting, northern blotting, and qPCR

15 cm dishes of NPM1-mScarlet expressing cells were seeded 24-48 hr prior to experiment such that cells were 70% confluent at time of harvest. Cells were washed once with ice cold PBS and then scraped into 1 mL ice cold lysis buffer (Pierce IP Lysis Buffer, ThermoFisher) supplemented with 0.5 mM TCEP and protease inhibitors (cOmplete Mini Tablets, Roche). Cells were sonicated at 30% power for 20 sec followed by ∼2 min on ice, three times in total. Cells were then spun at 16,000 x g at 4°C for 10 min to remove cell debris. An aliquot of lysate was removed for input samples, and then the remainder was split into two parts and each bound to 25 µL RFP-Trap resin (slurry, which was first washed with 1 mL of lysis buffer). Samples were bound for 1 hr at 4°C while rotating. Resin was washed twice with lysis buffer and then 4 times with wash buffer (25 mM Tris pH 7.4, 150 mM NaCl, 1mM EDTA, 0.05% NP-40, 0.5 mM TCEP). Samples for western blot were then eluted directly in 50 µL sample buffer (2% SDS, 10 mM EDTA, 10% glycerol, 0.1% bromophenol blue, 50 mM Tris, pH 7.4, 100 mM DTT). Samples for qPCR were eluted directly into 100 µL TRIzol reagent (ThermoFisher). Sample preparation then continued as described below.

### RT-qPCR

Cells were transfected in 24-well format, see siRNA transfection protocol above. After 72 hr of gene knockdown, cells were rinsed twice with ice-cold PBS (Gibco) and then scraped into 200 µL TRIzol Reagent (Invitrogen) and lysed by pipetting. Lysed cells were incubated for 5 min at room temperature. 40 µL chloroform was added and tubes were mixed by inversion followed by incubation for 2 minutes at room temperature. Samples were centrifuged for 15 min at 12,000 x g at 4°C. The aqueous phase was transferred to a new tube. RNA was extracted using Zymo RNA Clean and Concentrator kit according to manufacturer’s protocol. RNA was eluted in 15 µL RNase-free water (Ambion). cDNA was generated from 1 µg total RNA using iScript cDNA Synthesis Master Mix (Bio-Rad). cDNA was then diluted 1:20 for rRNA intermediates and 1:100 for mature rRNAs. RT-qPCR was performed using SYBR Green PCR Master Mix (Applied Bioystems) according to manufacturer’s protocol with a Bio-Rad CFX 96 Real Time thermal cycler. Reactions were run in technical triplicate and GAPDH was used as a housekeeping gene with the following primers:

GAPDH-F (5’-GTCTCCTCTGACTTCAACAGCG-3’)

GAPDH-R (5’-ACCACCCTGTTGCTGTAGCCAA-3’)

Primers used to amplify rRNA regions were as follows:

47S-F (5’-GAACGGTGGTGTGTCGTT-3’)

47S-R (5’-GCGTCTCGTCTCGTCTCACT-3’)

18S_5’-F (5’-GCCGCGCTCTACCTTACCTACCT-3’)

18S_5’-R (5’-CAGACATGCATGGCTTAATCTTTG-3’)

18S_3’-F (5’-AGTCGTAACAAGGTTTCCGTAGGT-3’)

18S_3’-R (5’-CCTCCGGGCTCCGTTAAT-3’)

5.8S_5’-F (5’-TACGACTCTTAGCGGTGGATCA-3’)

5.8S_5’-R (5’-TCACATTAATTCTCGCAGCTAGCT-3’)

5.8S_3’-F (5’-GAATTGCAGGACACATTGATCATC-3’)

5.8S_3’-R (5’-GGCAAGCGACGCTCAGA-3’)

18S-F (5’-CTGGATACCGCAGCTAGGAA-3’)

18S-R (5’-GAATTTCACCTCTAGCGGCG-3’)

5.8S-F (5’-ACTCGGCTCGTGCGTC -3’)

5.8S-R (5’-GCGACGCTCAGACAGG-3’)

28S-F (5’-CGGCGGGAGTAACTATGACT-3’)

28S-R (5’-GCTGTGGTTTCGCTGGATAG-5’)

Primers used for assessing gene depletions were as follows (designed using IDT PrimeTime):

MDN1 F: (5’-TTCGAGCACATTAAACAAGGC-3’)

MDN1 R: (5’-CCTCCTCTTCCTGATCTTTGC-3’)

NOG2 F: (5’-CCAATGAGAGCCACTTGT-3’)

NOG2 R: (5’-GATGGATTTGACCCTCACC-3’)

PHF5A F: (5’- ATTGTAAGGAGTCACCATCC-3’)

PHF5A R: (5’-GCGTTCATAGAAGAGGTCTGTC-3’)

CNOT1 F: (5’-CTGTCATACTGTTGCCACTGAT-3’)

CNOT1 R: (5’-CTCATACTCCCAACCTCTG-3’)

NSA2 F: (5’-TGAGACCCGAAAATTGAGAGC-3’)

NSA2 R: (5’-GGTAATCCAAACGGTATCCATAGC-3’)

RPF2 F: (5’-GTATGTTCTCACTTCACTGC-3’)

RPF2 R: (5’-ТСССАТСТСТТССААТТСААТСС-3’)

### Western blotting

Cells were washed twice with ice-cold PBS and then scraped into RIPA buffer (ThermoFisher) supplemented with protease inhibitors (cOmplete Mini Tablets, Roche). Cells were lysed by with a probe sonicator at 20% power in two 30-sec intervals on ice. Lysates were normalized by OD_260_ and denatured in sample buffer and boiled for 5 min at 95°C. Samples were run on a 4-20% Mini-PROTEAN TGX precast gel (Bio-Rad) and then transferred to nitrocellulose using a Bio-Rad TransBlot Turbo system. Membranes were blocked in 3% w/v BSA in TBST (50 mM Tris pH 8, 150 mM NaCl, 0.1% TWEEN-20) for 30 min and then incubated with primary antibodies overnight diluted at 1:1000 in antibody dilution buffer (3% BSA in TBST with 0.02% sodium azide). Membranes were washed three times in TBST and then incubated in 1:10,000 secondary antibody diluted in 3% BSA in TBST for 1-2 hr. Membranes were again washed three times in TBST and then incubated in SuperSignal West Pico Plus ECL substrate (ThermoFisher) and imaged on a Bio-Rad ChemiDOC MP Imaging system.

### RNA electrophoresis and northern blotting

RNA was harvested and purified as in IPs or from total cells for qPCR as described above. Eluted RNA was diluted 1:1 in formamide RNA loading buffer (1mM EDTA pH 8, 60mM Tricine, 60mM Triethanolamime, 95% formamide, 0.03% bromophenol blue, 7% formaldehyde, 30μg/mL Ethidium bromide), denatured at 70°C for five minutes and then snap cooled on ice. RNA was electrophoresed in a 1% agarose gel made with 1x tricine-triethanolamine (TT) buffer (30mM tricine, 30mM triethanolamine) and 7% formaldehyde for 2-8 hours at 180 V with recirculation. The gel was subjected to mild alkaline treatment (10 min 75mM NaOH, 10 min 0.5 M Tris pH 7.4, 1.5 M NaCl, 10 min in 10x SSC) and then transferred by capillary transfer to BrightStar-Plus positively charged nylon membrane overnight in 10x SSC. RNA was UV crosslinked to the membrane and then chemiluminescent northern blotting using biotinylated probes was performed as previously [92].

### Sucrose gradient fractionation

NPM1-mutant cell lines were plated 24-48 hours prior to fractionation and harvested at 70% confluency. Nuclear fractions were obtained as previously described by Pestov et al. and separated on a 10-30% linear sucrose gradient by ultracentrifugation with a Sorvall TH-641 rotor for 3 hours at 36,000 rpm and fractionated using a BioComp gradient fractionator. Fractions were pooled as described in text. For protein isolation, 1% BSA was added as a carrier and then protein was precipitated using TCA. For RNA isolation, 200 μL of the fraction was combined with 600 μL Trizol and purified as described above.

### Quantification of nucleolar RNA FISH and IF

Fixed-cell images were analyzed using custom pipelines developed in CellProfiler [93] (v.4.0.7). Briefly, for nucleolar RNA FISH, nucleoli were segmented using the NPM1-tagged channel using adaptive Otsu thresholding. There is a higher nucleoplasmic signal for FISH samples, and therefore, three threshold classes were used and the bottom two were classified as background. The object diameter range was set to 10-300 pixels, and objects outside this range were discarded. Nucleoli that touch the border of the image were discarded. Identified nucleoli were shrunk but 1 pixel to eliminate impacts of edge effects on intensity measurements. Intensity of identified nucleoli objects were measured for both the unscaled NPM1-tagged channel and the stained channel. For IF, the same pipeline was used, but nucleoli were thresholded using two class thresholding. A custom Python script was used for post-processing of CellProfiler data to bin images by field of view and experimental condition. In addition, nucleoli that had an NPM1 intensity level of less than 0.01 were eliminated to omit fields of view with no cells, which resulted in inappropriate identification of background noise.

### Partition coefficient calculations

Partition coefficients were determined manually by quantifying relative nucleolar (I^dense^) and nucleoplasmic (I^dilute^) NPM1 intensities from pre-bleach (time point 0) images acquired by HiT-FRAP, as follows: Nucleolar signal was determined by measuring the average intensity of a 10 x 10 pixel box in the most intense region of nucleolar signal (5 x 5 pixels were used when nucleoli were small). Nucleoplasmic signal was determined by measuring the average intensity of a 10 x 10 pixel box that was manually chosen in the same cell as the measured nucleolus. Background was determined by averaging the average intensity across 5 randomly chosen 10 x 10 pixel locations. Background was subtracted, and then the ratio of nucleolar to nucleoplasmic signals were calculated to determine partition coefficient (K = I^dense^/I^dilute^, where K = partition coefficient).

### RNA Helicase phylogenetic analysis

Full-length, reference protein sequences for indicated DEAD-box proteins were retrieved from NCBI and aligned using MUSCLE^101^. A maximum-likelihood tree was constructed using MEGA^102^ using default options and the JTT model.

### Statistical analysis

Software used for statistical analysis of screen-related data is listed in subsection of methods above. Otherwise, all other statistical tests were performed in Prism 8 as indicated in figure legends.

### Use of AI

Portions of the manuscript text were revised with the assistance of Claude, Sonnet 4.6. All content was reviewed by authors.

## Notes

### Competing Interest Statement

The authors have declared no competing interest.

