## Supplemental Text and Figures for "Nucleolar dynamics are determined by the ordered assembly of the ribosome"

**Supplementary Figures for Sheu-Gruttadauria et al., "Nucleolar dynamics are determined by the ordered assembly of the ribosome"**

### Supplementary Text

*The LSU phenotypic group can be defined by two phenotypically and functionally distinct subclusters*

The LSU phenotypic group that clustered according to gene function and PCA space could be further divided into two subgroups, LSU1 and LSU2, by z-score hierarchical clustering (Fig. 3D). These groups both exhibit an increase in nucleolar size, but LSU1 shows a more notable increase in area while LSU2 shows a more irregular shape and a substantial decrease in intensity and $t_{1/2}$ (Fig. S8A). While both factors are associated with late LSU maturation steps, LSU1 is enriched for factors that associate during earlier stages of LSU assembly (States A-F), while LSU2 is composed primarily of factors that bind at later stages (States G/H) (Fig. S-T B and C). Indeed, the most striking feature of LSU2 is the strong enrichment for factors involved in maturation of the central protuberance (CP), rotation of the L1 stalk, and rearrangement of the ITS2 foot for subsequent cleavage and processing (Fig. S8B and C). These results suggest that we may have identified phenotypically coherent and distinct groups associated with different stages in pre-LSU maturation.

To further explore differences between LSU1 and LSU2, we assessed rRNA intermediate composition for the strong hit NSA2 in the LSU1 cluster, as performed for LSU2 hits RPF2 and NOG2 (see main text). We saw a significant nucleolar depletion of the 5′ETS and ITS1 and accumulation of ITS2, with a reduction of the mature 28S by qPCR (Fig. S8D). These results suggest that like LSU2, stalled pre-LSU complexes are accumulating in the nucleolus upon depletion of LSU1 genes. We also performed IF for NSA2 to investigate what LSU precursors might accumulate. Unlike LSU2 genes, we found a strong enrichment for NOP2 and no change in NOP53 upon depletion of NSA2 (Fig. S8E), which suggests that states upstream of G and H may accumulate in the LSU1 phenotypic cluster. We also observed a modest increase in nucleolar NOG2, however. Therefore, it is possible that some of these upstream intermediates can still recruit downstream factors, in keeping with previous studies in yeast that demonstrate NOG2 can associate with earlier intermediates[64]. These results may suggest that the LSU1 phenotype may result from accumulation of earlier stalled intermediates than LSU2. However, future work will be necessary to more clearly distinguish between abortive nucleolar intermediates in these two clusters.

*Replication of major phenotypes in p53-WT hTERT-RPE cells*

To assess whether the nucleolar biophysical states we identified in HeLa cells reflect general biological responses to ribosome biogenesis disruption, we depleted representative hits from each phenotypic cluster in hTERT-RPE cells, a diploid, p53-proficient line, transduced with lentiviral NPM1-mScarlet, and assessed nucleolar morphology in the polyclonal cell line (Fig. S6C). We note that baseline nucleolar size was smaller in RPE than HeLa cells, consistent with lower baseline ribosome biogenesis activity reported in non-transformed cells. We found that depletion of SKIV2L2 from the "RNA exosome" cluster shows a different phenotype than seen in HeLa cells, where we observe less size difference and a marked decrease in eccentricity, which may reflect a p53 or cell type specific response. Depletion of the “Other” cluster gene PHF5A results in a milder though qualitatively similar phenotype as seen in HeLa cells, with nucleolar rounding and an increase in NPM1 intensity. Depletion of “LSU”-associated hits in RPE cells very robustly replicated most of the nucleolar features we observed in HeLa, which suggest that these are likely generalizable responses to LSU disruption.

*Further characterization of “Other” phenotypic cluster*

To confirm “Other” phenotypes, we used inhibitors of the two most represented pathways, NMD and pre-mRNA splicing. Treating cells for 24 hours with the small molecule SMG1i, which prevents activation of the core NMD RNA helicase UPF1[65], phenocopied genetic ablation of NMD factors (Fig. S9C). Similarly, treating cells with an inhibitor of the SF3b complex pladienolide B[66] (PladB) for 24 hours recapitulated depletion of splicing-associated factors (Fig. S9D). Notably, although defects in dynamics were only detectable after 24 hours of treatment, nucleolar rounding occurred on shorter (0-4hr) time scales, which suggests an acute impact of splicing inhibition on nucleolar assembly (Fig. S9E). Together, these observations validate our screen results. However, we note that the NMD-associated phenotype was less robust in the mScarlet cell line (Fig. S2C) and therefore, we chose to focus further characterization on more reproducible hits.

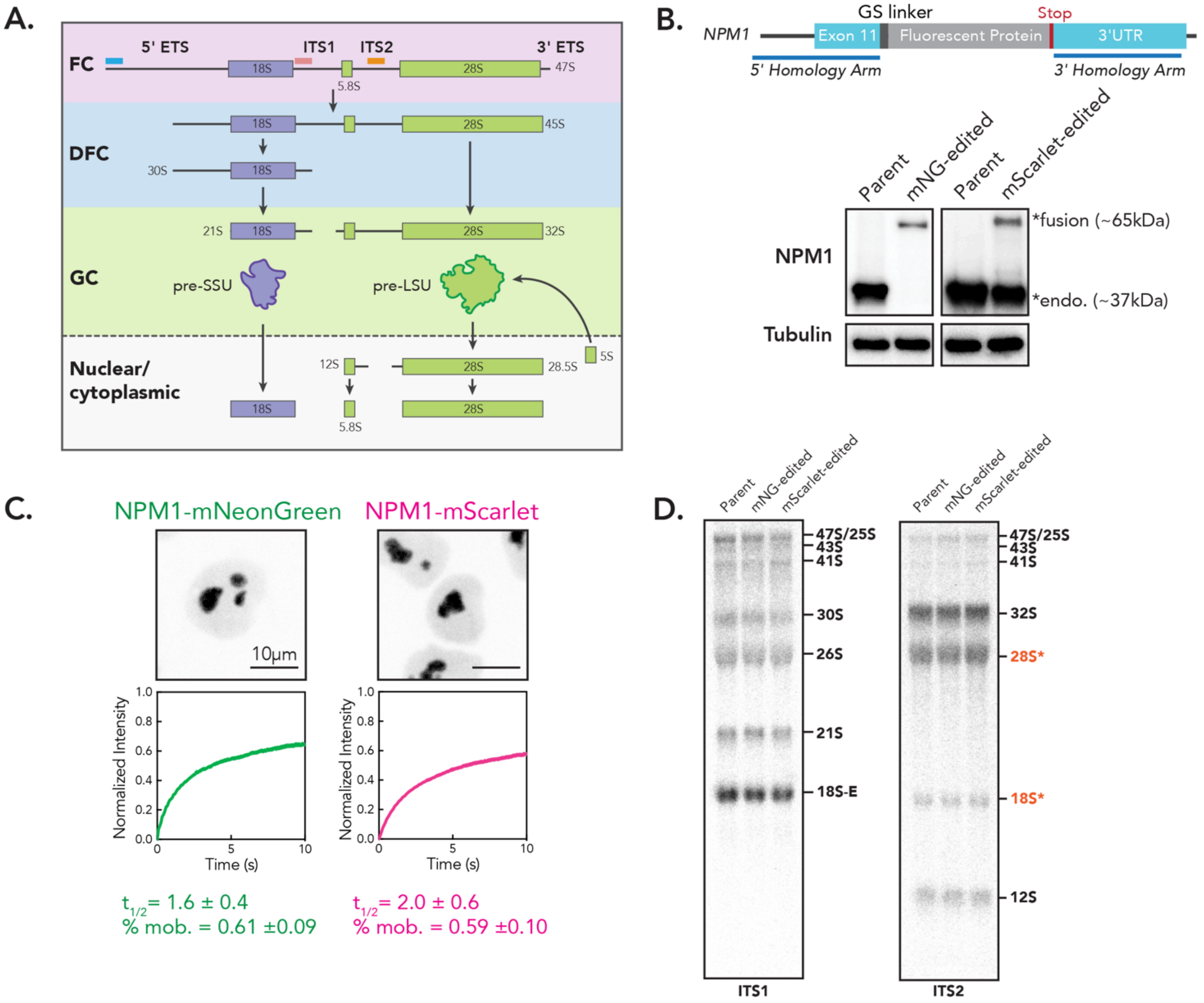


**Figure S1: Schematic of pre-rRNA processing and NPM1 reporter cell line generation and characterization.** (A) Schematic showing subset of rRNA processing steps in humans, with approximate localizations indicated. (B) Schematic for endogenous insertion of fluorescent protein tag at C-terminus of NPM1. Western blots for both mNeonGreen and mScarlet edited cell lines. Fusion protein and endogenous protein indicated. (C) Images and FRAP curves for mNeonGreen and mScarlet tagged lines. Scale bar is 10 µm. FRAP curves for n = 250 nucleoli, error bars are 95% CI. $t_{1/2}$ and mobile fraction shown, mean ± SD. (D) Northern blot for cell lines used in this manuscript. ITS1 and ITS2 blots shown. Nonspecific signal from 28S and 18S indicated.

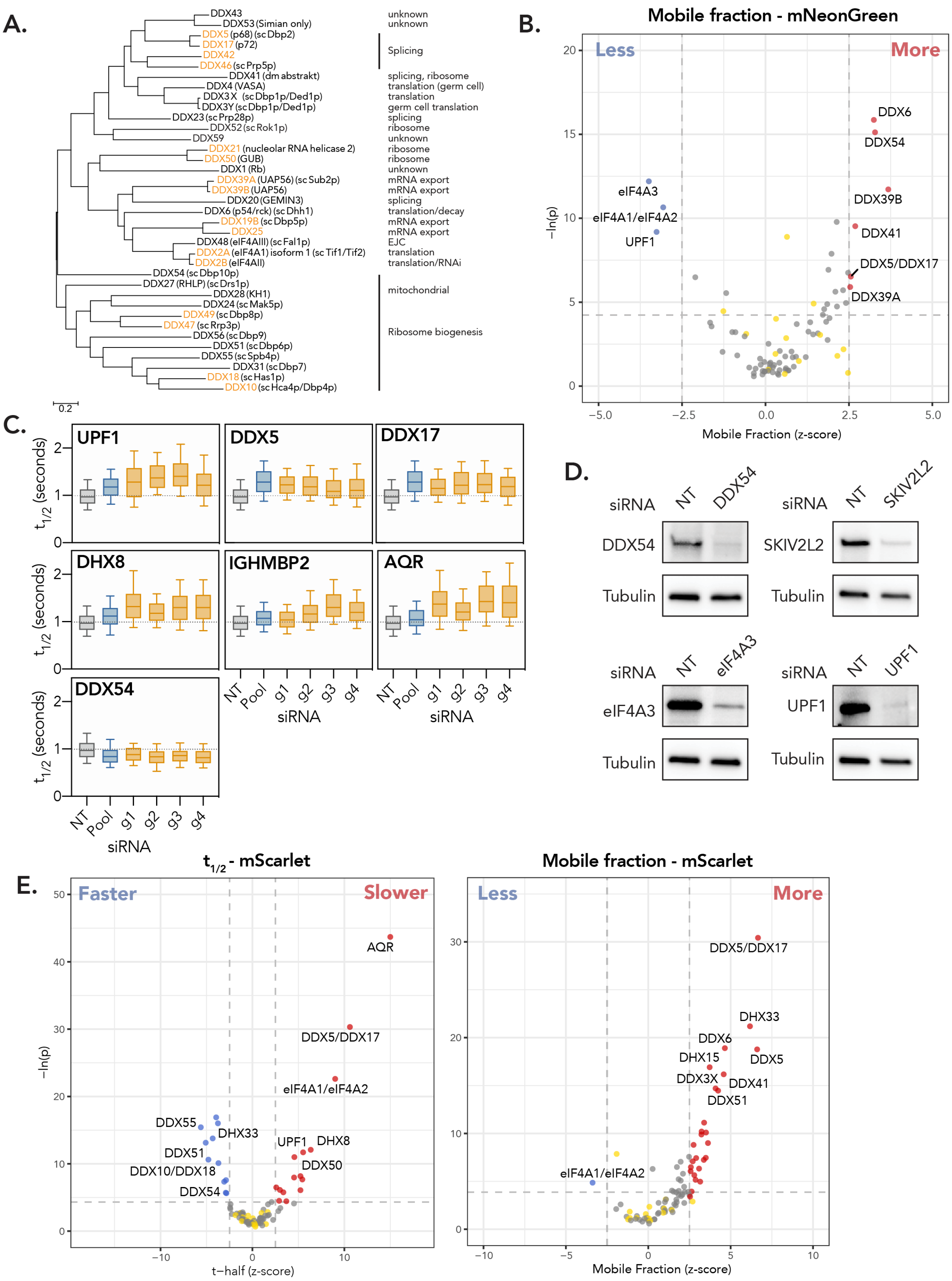
A.
DDX43
DDX53 (Simian only)
DDX5 (p68) (sc Dbp2)
DDX17 (p72)
DDX42
DDX46 (sc Prp5p)
DDX41 (dm abstrakt)
DDX4 (VASA)
DDX3X (sc Dbp1p/Ded1p)
DDX3Y (sc Dbp1p/Ded1p)
DDX23 (sc Prp28p)
DDX52 (sc Rok1p)
DDX59
DDX21 (nucleolar RNA helicase 2)
DDX50 (GUB)
DDX1 (Rb)
DDX39A (UAP56) (sc Sub2p)
DDX39B (UAP56)
DDX20 (GEMIN3)
DDX6 (p54/rck) (sc Dhh1)
DDX19B (sc Dbp5p)
DDX25
DDX48 (eIF4AIII) (sc Fal1p)
DDX2A (eIF4A1) isoform 1 (sc Tif1/Tif2)
DDX2B (eIF4AII)
DDX54 (sc Dbp10p)
DDX27 (RHLP) (sc Drs1p)
DDX28 (KH1)
DDX24 (sc Mak5p)
DDX49 (sc Dbp8p)
DDX47 (sc Rrp3p)
DDX56 (sc Dbp9)
DDX51 (sc Dbp6p)
DDX55 (sc Spb4p)
DDX31 (sc Dbp7)
DDX18 (sc Has1p)
DDX10 (sc Hca4p/Dbp4p)
0.2
unknown
unknown
Splicing
splicing, ribosome
translation (germ cell)
translation
germ cell translation
splicing
ribosome
unknown
ribosome
ribosome
unknown
mRNA export
mRNA export
splicing
translation/decay
mRNA export
mRNA export
EJC
translation
translation/RNAi
mitochondrial
Ribosome biogenesis
B.
Mobile fraction - mNeonGreen
Less
More
DDX6
DDX54
eIF4A3
eIF4A1/eIF4A2
UPF1
DDX39B
DDX41
DDX5/DDX17
DDX39A
−ln(p)
Mobile Fraction (z-score)
C.
UPF1
DDX5
DDX17
DHX8
IGHMBP2
AQR
DDX54
t1/2 (seconds)
NT
Pool
g1
g2
g3
g4
siRNA
D.
siRNA
NT
DDX54
SKIV2L2
eIF4A3
UPF1
Tubulin
E.
t1/2 - mScarlet
Faster
Slower
AQR
DDX5/DDX17
eIF4A1/eIF4A2
DDX55
DHX33
DDX51
DDX10/DDX18
DDX54
UPF1
DHX8
DDX50
−ln(p)
t−half (z-score)
Mobile fraction - mScarlet
Less
More
DDX5/DDX17
DHX33
DDX6
DHX15
DDX5
DDX3X
DDX41
DDX51
eIF4A1/eIF4A2
Mobile Fraction (z-score)

**Figure S2: HiT-FRAP dynamics screen validation.** (A) Maximum-likelihood phylogenetic tree for human DEAD-box helicases, with parenthetical orthologs and putative functions. Helicase pairs depleted in primary screen shown in orange, except for DDX19A/DDX19B. Scale bar represents mean number of nucleotide substitutions per site. (B) Volcano plot for mobile fraction for primary RNA helicase screen, highlighting helicase hits that resulted in increased (red) or decreased (blue) mobile fraction relative to non-targeting control cells (yellow; FDR $< 0.05$ and z-cutoff of $\pm$ 2.5 shown as dotted lines, see methods). Non-significant gene targets shown in gray. x-axis plots robust z-score for mobile fraction (see methods). (C) Depool validation of siRNA pool gene depletion strategy. $t_{1/2}$ shown for NT (gray) and pool (blue) of indicated gene depletion. Individual guides shown in yellow. Values have been normalized to pool or single non-targeting guides, respectively. Error bars are 10-90 percentile. (D) Western blot for select hits to validate knockdown. (E) Volcano plot for $t_{1/2}$ (left) and mobile fraction (right) for primary RNA helicase screen performed in NPM1-mScarlet reporter cell line, highlighting helicase hits that resulted in increased (red, "slower" and "more" for $t_{1/2}$ and mobile fraction, respectively) or decreased (blue, "faster" and "less" for $t_{1/2}$ and mobile fraction, respectively) $t_{1/2}$ or mobile fraction relative to non-targeting control cells (yellow; FDR $< 0.05$ and z-cutoff of $\pm$ 2.5 shown as dotted lines, see methods). Non-significant gene targets shown in gray. x-axis plots robust z-score for $t_{1/2}$ (see methods).

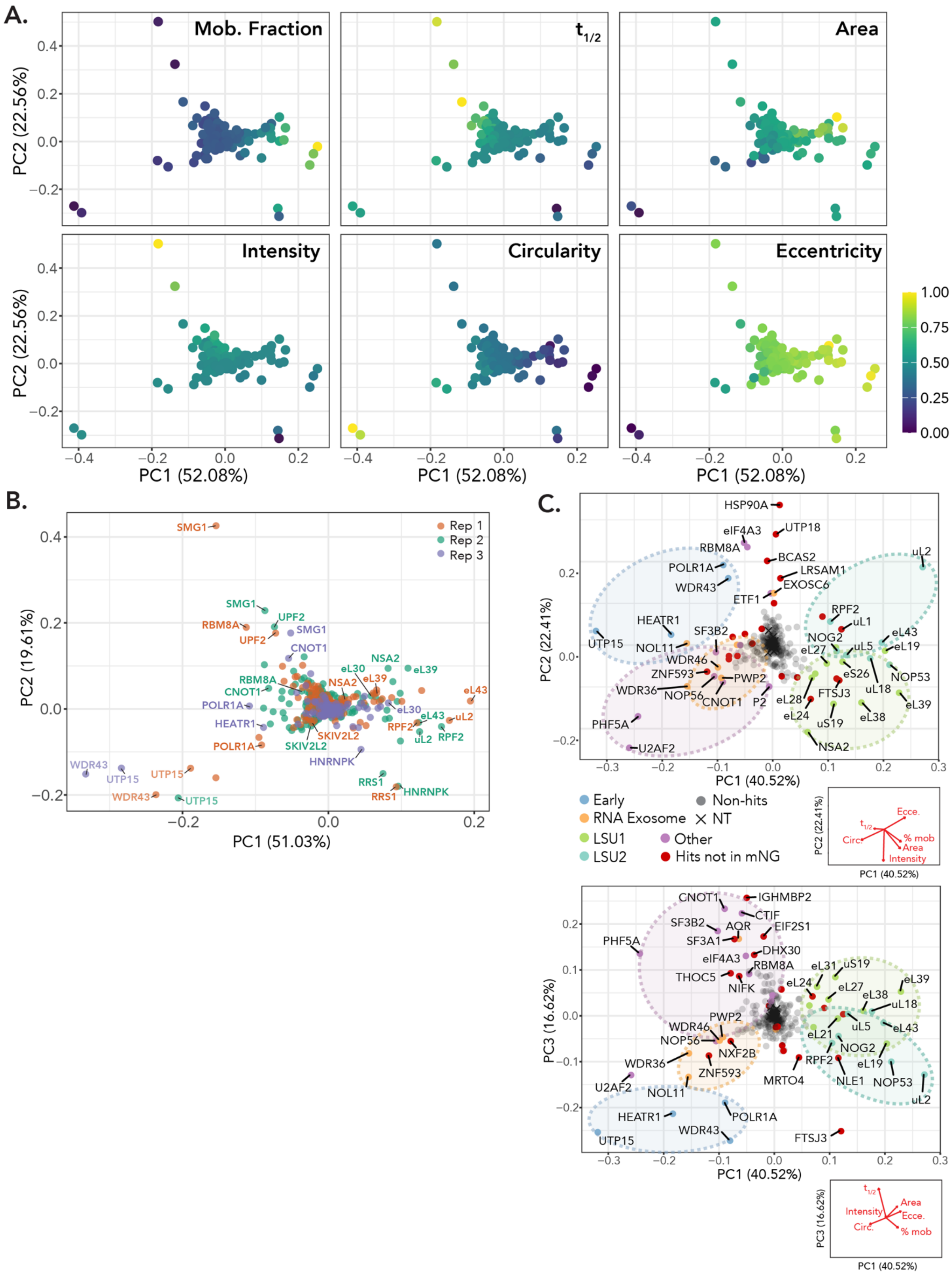
A.
Mob. Fraction
t1/2
Area
Intensity
Circularity
Eccentricity
PC2 (22.56%)
PC1 (52.08%)
B.
Rep 1
Rep 2
Rep 3
PC2 (19.61%)
PC1 (51.03%)
C.
PC2 (22.41%)
PC1 (40.52%)
Early
RNA Exosome
LSU1
LSU2
Non-hits
NT
Other
Hits not in mNG
PC3 (16.62%)

**Figure S3: Phenotypic clustering features and validation.** (A) PCA colored by six nucleolar features used in analysis. Viridis scale used, scale bar indicated. (B) Three biological replicates (in NPM1-mNeonGreen) cell line shown on same PCA. Select hits that fall in similar regions across 2 out of 3 replicates indicated. (C) PCA of replicated screens in NPM1-mScarlet cell line. Colored by manually annotated function. PC2 vs. PC1 and PC3 vs. PC1 shown. PCA loadings shown below.

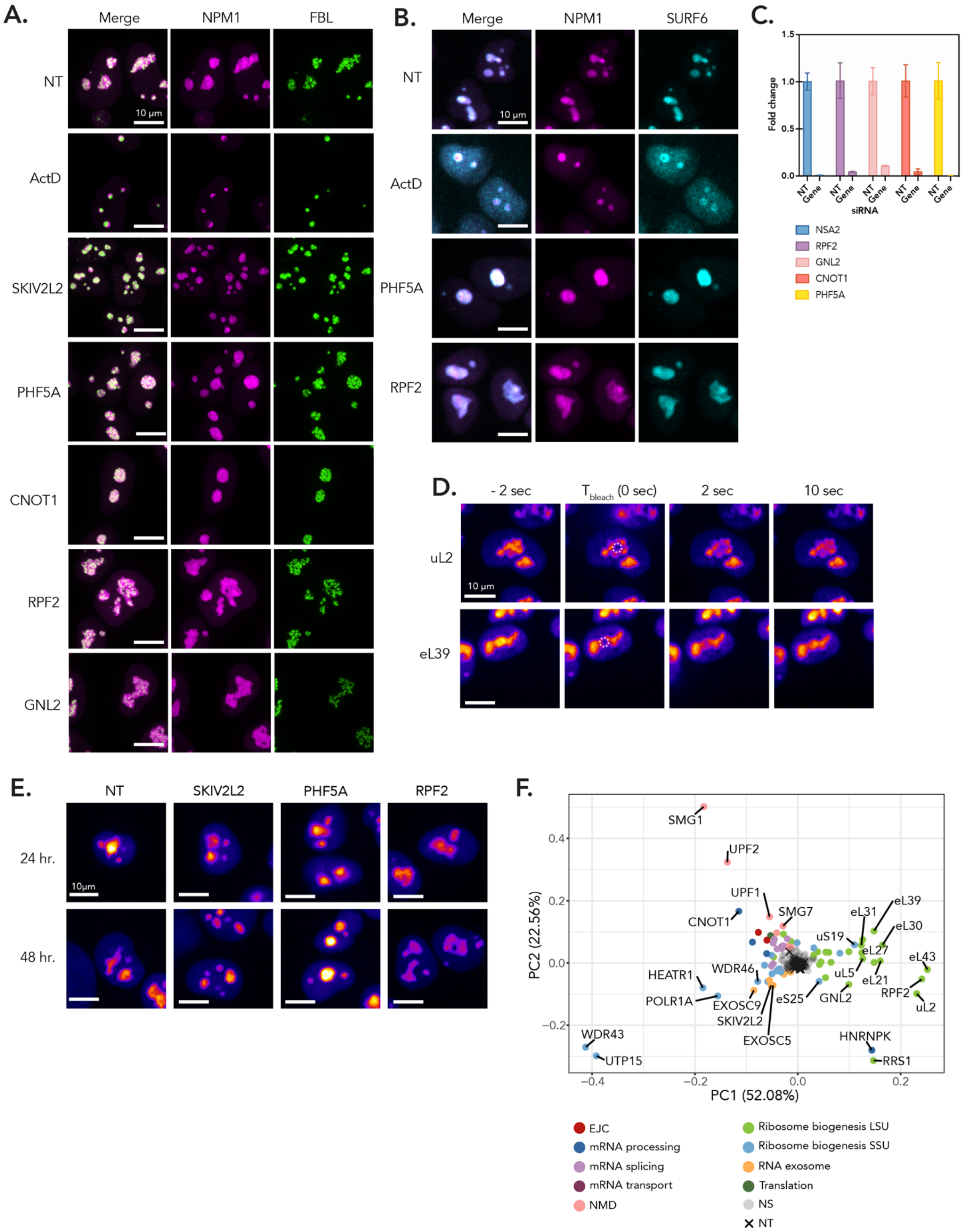
A.
Merge
NPM1
FBL
NT
ActD
SKIV2L2
PHF5A
CNOT1
RPF2
GNL2
10 μm
B.
Merge
NPM1
SURF6
NT
ActD
PHF5A
RPF2
10 μm
C.
Fold change
1.5
1.0
0.5
0.0
NT
Gene
siRNA
NSA2
RPF2
GNL2
CNOT1
PHF5A
D.
- 2 sec
T bleach (0 sec)
2 sec
10 sec
uL2
eL39
10 μm
E.
NT
SKIV2L2
PHF5A
RPF2
24 hr.
48 hr.
10μm
F.
PC2 (22.56%)
PC1 (52.08%)
0.4
0.2
0.0
−0.2
−0.4
−0.2
0.0
0.2
SMG1
UPF2
UPF1
SMG7
CNOT1
eL31
eL39
eL30
uS19
eL27
eL43
uL5
eL21
RPF2
uL2
WDR46
HEATR1
POLR1A
EXOSC9
eS25
GNL2
SKIV2L2
EXOSC5
WDR43
UTP15
HNRNPK
RRS1
EJC
mRNA processing
mRNA splicing
mRNA transport
NMD
Ribosome biogenesis LSU
Ribosome biogenesis SSU
RNA exosome
Translation
NS
NT

**Figure S4: Phenotypic clustering and validation.** (A) Dual color cell line (GC: NPM1-mScarlet, DFC: FBL-mScarlet, endogenously tagged) images for representative hits from phenotypic groups. Maximum intensity z-projection. Scale bar is 10 $\mu$m. (B) Immunofluorescence staining for GC marker SURF6 in representative hits from Early (ActD), Other (PHF5A), and LSU (RPF2) clusters. Scale bar is 10 $\mu$m. (C) qPCR showing fold change gene expression for select hits in each phenotypic cluster. NT = non-targeting control, Gene = siRNA against gene indicated. Internally normalized to GAPDH. Error bars are SD, n = 3. (D) Bleaching time course for disrupted LSU hits uL2 and eL39 showing bleach points target the nucleolar interior. Scale bar is 10 $\mu$m. (E) Depletion time course for representative hits from RNA Exosome (SKIV2L2), Other (PHF5A), and LSU (RPF2) phenotypic groups. Representative images of nucleolar phenotypes upon 24 hours and 48 hours post-siRNA transfection shown. Images scaled equally and colored with mpl-inferno LUT to show NPM1 intensity. Scale bar is 10 $\mu$m. (F) NPM1-mNeonGreen screen PCA shown colored by manually annotated function.

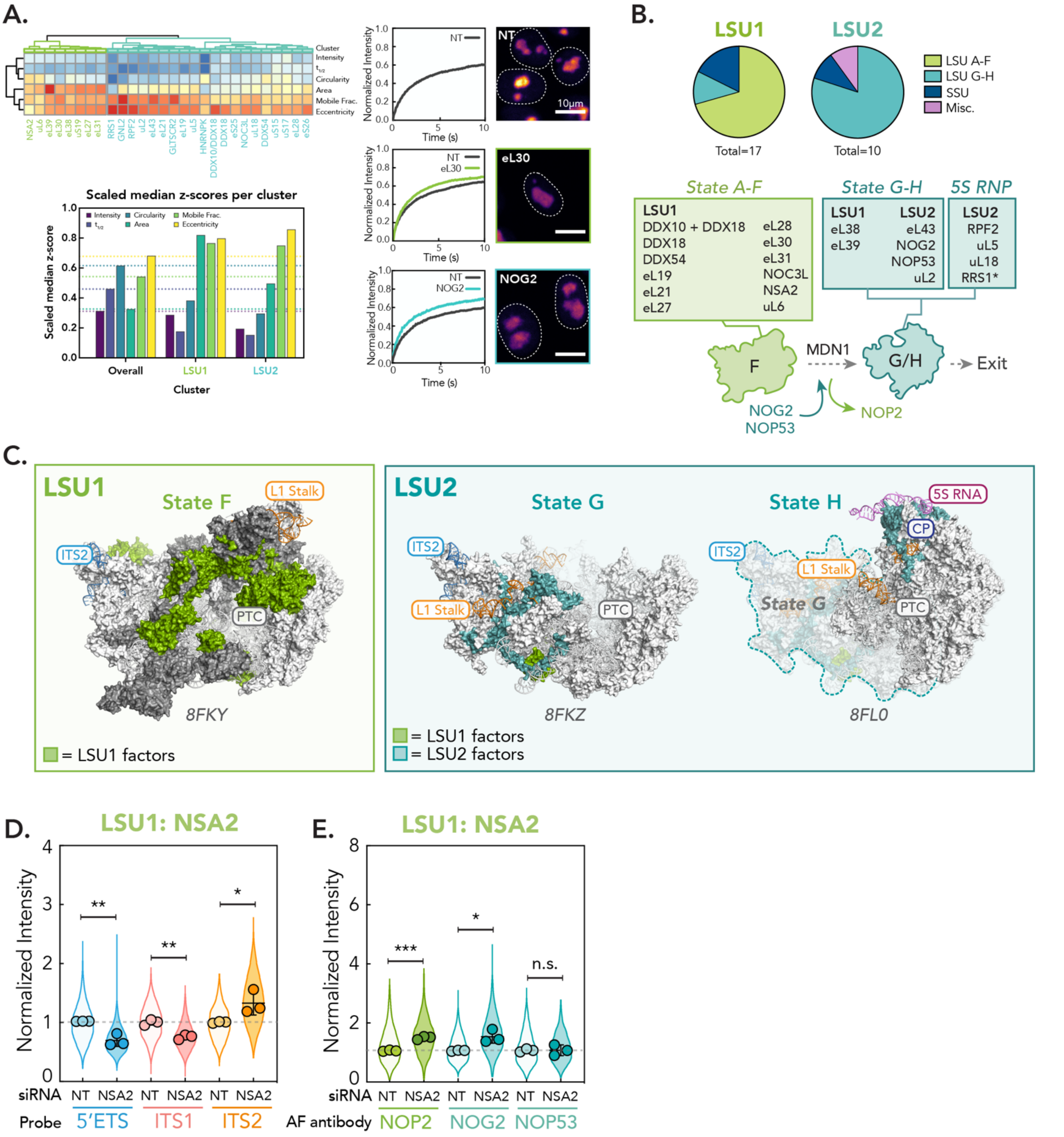


**Figure S5: Phenotypic clustering reveals sub-clusters associated with functionally coherent genes in late-LSU assembly.** (A) Cropped heatmap from Fig. 3D showing LSU cluster split into two phenotypic clusters by hierarchical clustering (LSU1 and LSU2, respectively). Median feature z-scores for clusters shown in bar graph, where dotted line represents median z-scores across all hits that went into the analysis. Representative images and FRAP curves for of hits from LSU1 and LSU2 shown. Scale bar is 10 μm. Note, NT and

NOG2 figure panels are replicated from Fig. 3F. (B) Mapping genes in LSU1 and LSU2 phenotypic clusters onto LSU intermediates and schematic of State F to G transition. * = RRS1 associates in State A-F but is critical in 5S RNP recruitment. (C) LSU1 (green) and LSU2 (teal) hits mapped onto pre-LSU intermediate structures. Major landmarks noted for reference. On State F, additional factors that are part of protein interaction network removed by MDN1 noted in dark gray. State H, which was a partial reconstruction, is overlayed on State G (transparent). PDB IDs: State F, 8FKY; State G, 8FKZ; State H, 8FL0. (D) rRNA FISH for representative hit in LSU1 (NSA2), as in Fig. 4B. (E) Nucleolar immunofluorescence of LSU intermediate markers for LSU1 representative gene hit NSA2. Three biological replicates were performed (n = 600 nucleoli per replicate, spread of individual nucleoli shown as violin, error bars are SD). Dotted line shows non-targeting control level. p-values calculated using two-tailed unpaired t-test between biological replicates. n.s. = not significant, * $p < 0.05$, ** $p<0.01$, *** $p < 0.001$.

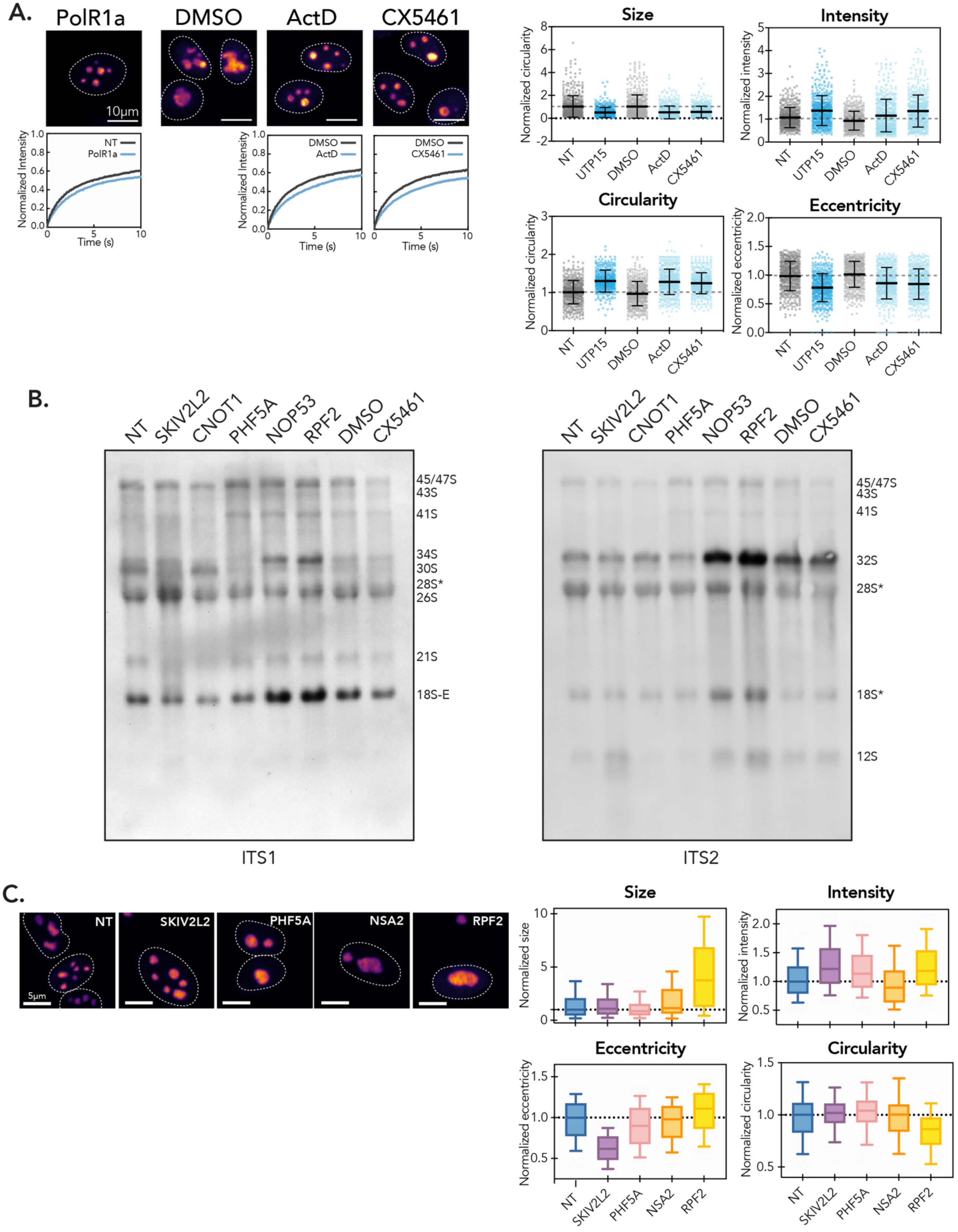
A.
PolR1a
DMSO
ActD
CX5461
10μm
Normalized Intensity
NT
PolR1a
Time (s)
ActD
Size
Intensity
Circularity
Eccentricity
Normalized circularity
Normalized intensity
Normalized eccentricity
NT
UTP15
DMSO
ActD
CX5461
B.
NT
SKIV2L2
CNOT1
PHF5A
NOP53
RPF2
DMSO
CX5461
45/47S
43S
41S
34S
30S
28S*
26S
21S
18S-E
32S
18S*
12S
ITS1
ITS2
C.
NT
SKIV2L2
PHF5A
NSA2
RPF2
5μm
Normalized size
Normalized intensity
Normalized eccentricity
Normalized circularity

**Figure S6: Further validation of phenotypic clusters.** (A) Comparison between PolR1a knockdown (Early cluster) phenotype and treatment of cells with Pol I inhibitors (0.04 $\mu$g/mL ActD and 500 nM CX5461 for 2 hr). Images are scaled equally and colored with mpl-inferno LUT to show relative intensity differences of NPM1. Dotted line shows nuclear boundary. Scale bar is 10 $\mu$m. FRAP curves for NT/vehicle control vs. gene depletion/drug shown (n = 250 nucleoli, error bars are 95% CI). Right, additional feature parameters for Pol1 inhibition as compared to Early cluster gene UTP15 depletion. Points are individual nucleoli, n = 500. Error bars are mean ± SD. Normalized to appropriate control to account for imaging differences. Dotted line indicates control level. (B) ITS1 and ITS2 northern blot for representative hits across phenotypic clusters. * = Nonspecific 28S and 18S. (C) Representative images of hit knockdowns in hTERT-RPE1 cells lentivirally expressing NPM1-mScarlet. Key morphological features shown below, normalized to NT. Box and whisker shows median and 10-90 percentile. n = 500 nucleoli.

(A) Comparison between PolR1a knockdown (Early cluster) phenotype and treatment of cells with Pol I inhibitors (0.04 $\mu$g/mL ActD and 500 nM CX5461 for 2 hr). Images are scaled equally and colored with mpl-inferno LUT to show relative intensity differences of NPM1. Dotted line shows nuclear boundary. FRAP curves for NT/vehicle control vs. gene depletion/drug shown (n = 250 nucleoli, error bars are 95% CI).

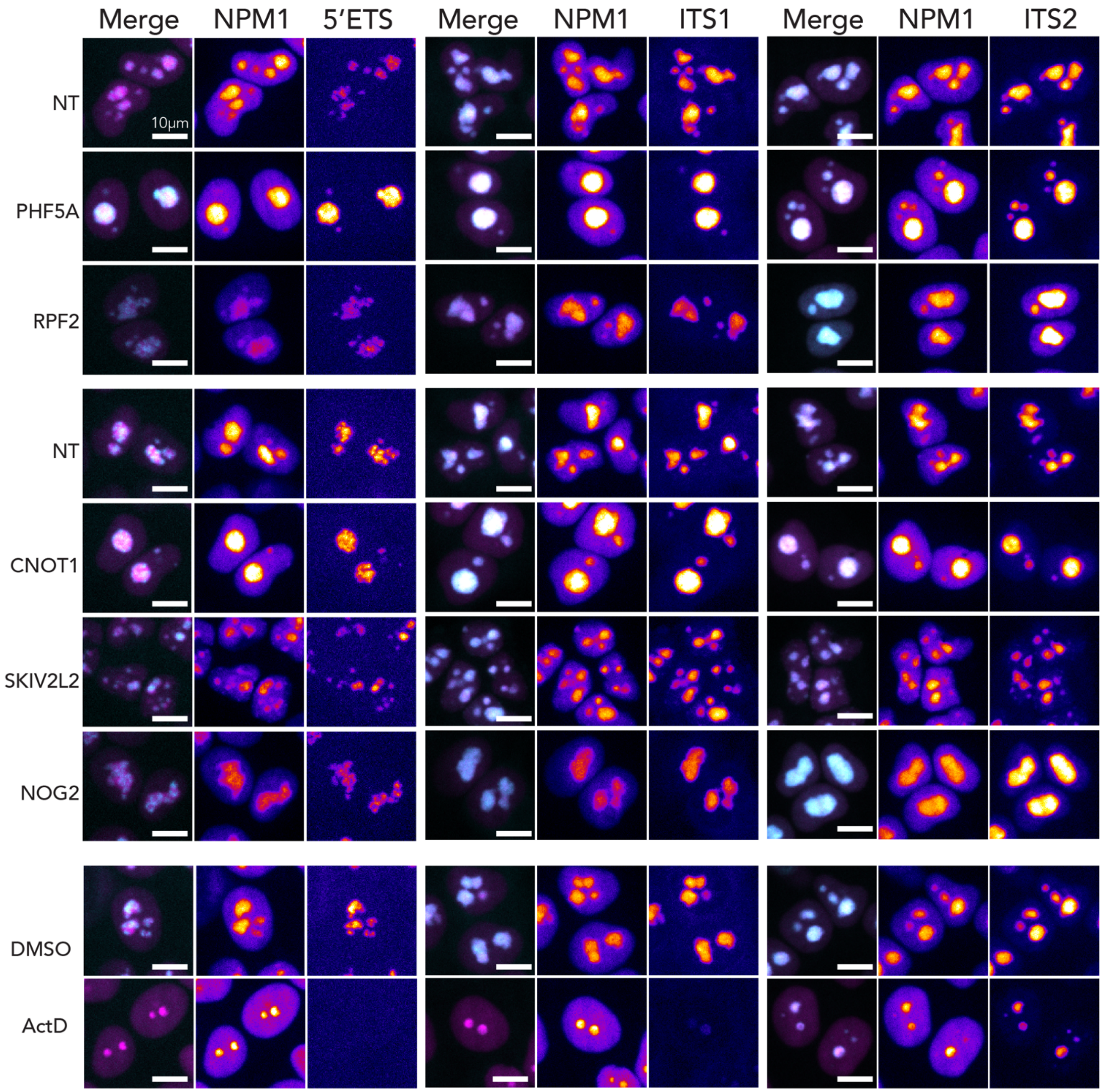


**Figure S7: Representative images for nucleolar rRNA FISH.** Merge shows pseudocolor NPM1 (magenta) and rRNA (cyan). NPM1 and rRNA panels colored by Fire-LUT for each marker, respectively, to show intensity differences. Internal controls shown for each group of gene depletions. All images scaled equally with respect to internal control. Scale bar is 10 μm.

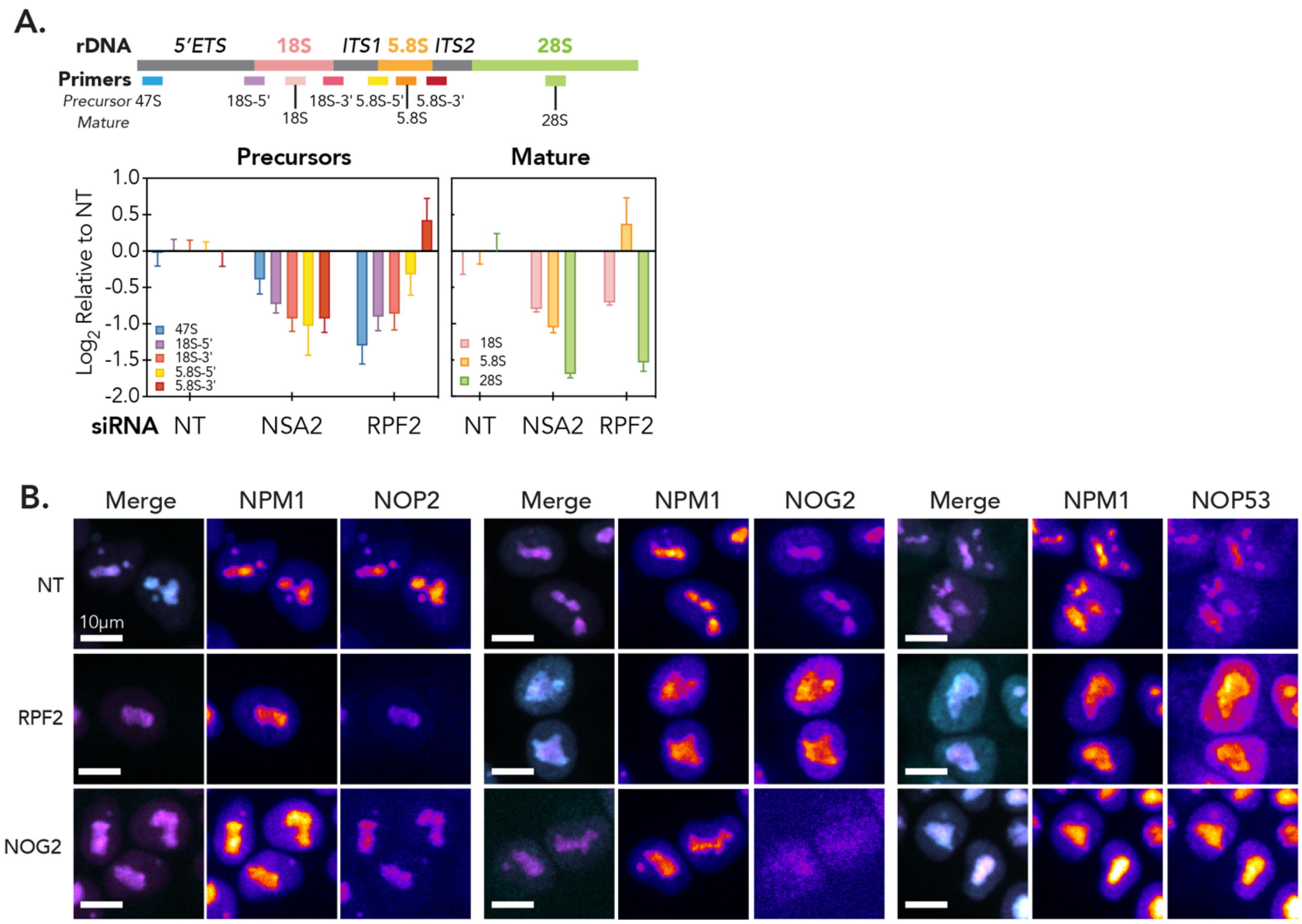


**Figure S8: Additional characterization of LSU-associated phenotypic clusters.** (A) qPCR to assess global rRNA precursor and mature rRNA levels for LSU1 (NSA2) and LSU2 (RPF2) genes (See supplementary text for discussion of LSU sub-clusters). Schematic shows positions of primers used. Error bars are SD of three biological replicates. (B) Representative immunofluorescence images shown for LSU hits. Merge shows pseudocolored markers NPM1 (magenta) and assembly factor (cyan). NPM1 and assembly factor panels colored by Fire-LUT for each marker, respectively, to show intensity differences. Internal controls shown for each group of gene depletions. All images scaled equally with respect to internal control. Scale bar is 10 μm.

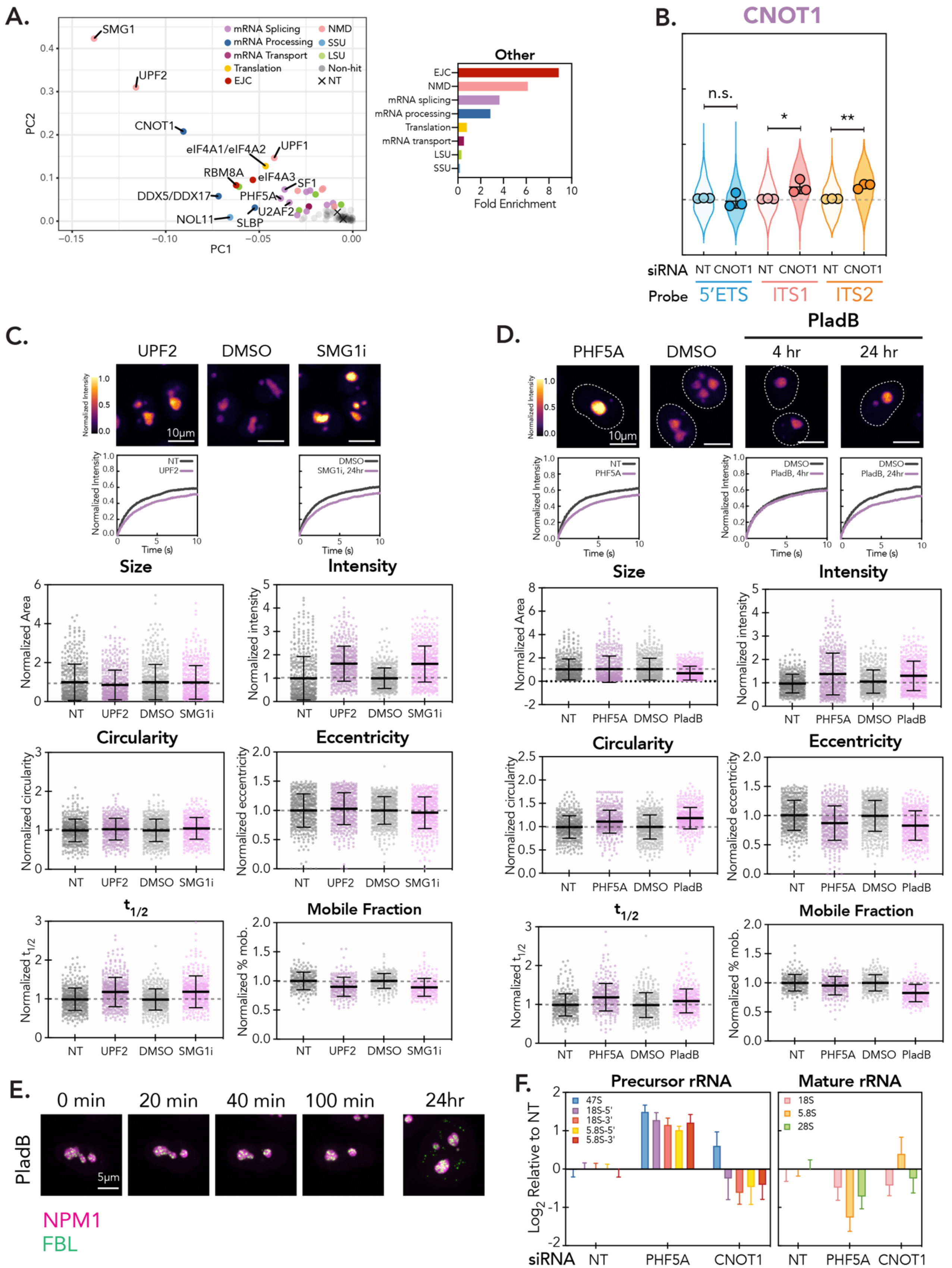
A.
B.
CNOT1
C.
D.
E.
F.
SMG1
UPF2
CNOT1
eIF4A1/eIF4A2
UPF1
RBM8A
eIF4A3
SF1
DDX5/DDX17
PHF5A
U2AF2
NOL11
SLBP
mRNA Splicing
mRNA Processing
mRNA Transport
Translation
EJC
NMD
SSU
LSU
Non-hit
NT
PC1
PC2
Other
Fold Enrichment
n.s.
siRNA
Probe
5′ETS
ITS1
ITS2
UPF2
DMSO
SMG1i
PHF5A
PladB
4 hr
24 hr
Normalized Intensity
Time (s)
Size
Intensity
Circularity
Eccentricity
$t_{1/2}$
Mobile Fraction
Normalized Area
Normalized intensity
Normalized circularity
Normalized eccentricity
Normalized $t_{1/2}$
Normalized % mob.
0 min
20 min
40 min
100 min
24hr
NPM1
FBL
Precursor rRNA
Mature rRNA
$Log_2$ Relative to NT
47S
18S-5'
18S-3'
5.8S-5'
5.8S-3'
18S
5.8S
28S

**Figure S9: Additional characterization of “Other” phenotypic cluster.** (A) “Other” phenotypic cluster shown as zoom of PCA in Fig. 3C. Fold enrichment for functional groups in “Other” cluster shown on right. (B) rRNA FISH for representative hit CNOT1 as in Fig. 4B. (C) Comparison between “Other” hit UPF2 and treatment with NMD inhibitor SMG1i (300 nM for 24 hr). Images are scaled equally and colored with mpl-inferno LUT to show relative intensity differences. Scale bar is 10 μm. FRAP curves for NT/vehicle vs. gene depletion/SMG1i shown (n = 250 nucleoli, error bars are 95% CI). Below, additional feature parameters for SMG1i treatment as compared to Other cluster gene UPF2 knockdown. Points are individual nucleoli, n = 500. Error bars are mean ± SD. Normalized to appropriate control to account for imaging differences. (D) Comparison between “Other” representative hit PHF5A and treatment with splicing inhibitor PladB (10 nM for times indicated). Images are scaled equally and colored with mpl-inferno LUT to show relative intensity differences of NPM1. Scale bar is 10 μm. FRAP curves for NT/vehicle vs. gene depletion/PladB shown (n = 250 nucleoli, error bars are 95% CI). Below, additional feature parameters for PladB inhibition (24 hr) as compared to Other cluster gene PHF5A knockdown. Points are individual nucleoli, n = 500. Error bars are mean ± SD. Normalized to appropriate control to account for imaging differences. (E) Time course of PladB treatment (10 nM) using dual color cell line shown as merge of NPM1 (magenta) and FBL (green). Scale bar is 5 μm. (F) qPCR to assess global rRNA precursor and mature rRNA levels for Other genes PHF5A and CNOT1.

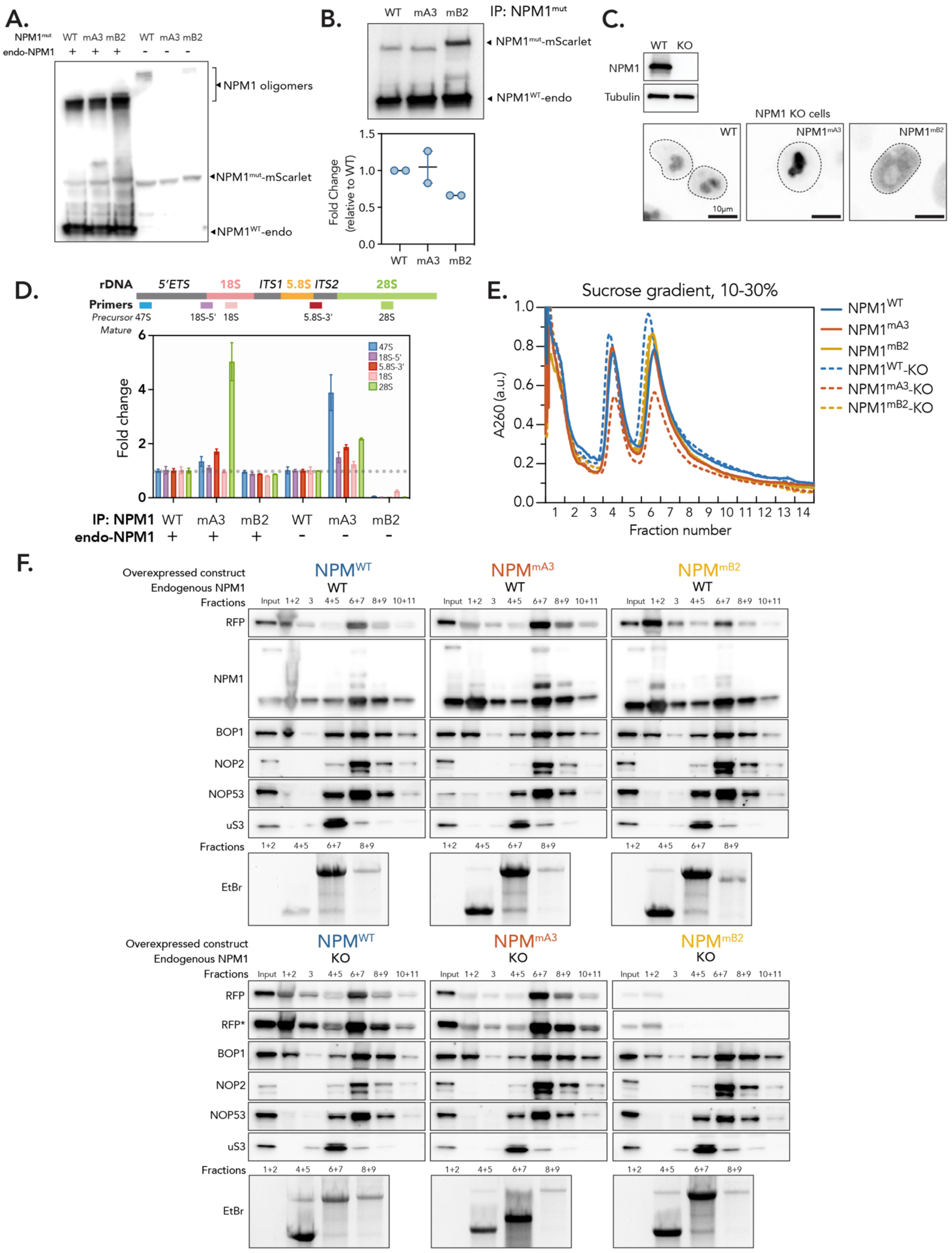
A.
NPM1mut WT mA3 mB2 WT mA3 mB2
endo-NPM1 + + + - - -
NPM1 oligomers
NPM1mut-mScarlet
NPM1WT-endo
B.
WT mA3 mB2
IP: NPM1mut
NPM1mut-mScarlet
NPM1WT-endo
Fold Change (relative to WT)
C.
WT KO
NPM1
Tubulin
NPM1 KO cells
WT
NPM1mA3
NPM1mB2
10µm
D.
rDNA
5'ETS 18S ITS1 5.8S ITS2 28S
Primers
Precursor 47S
Mature
18S-5' 18S 5.8S-3' 28S
Fold change
47S
18S-5'
5.8S-3'
18S
28S
IP: NPM1 WT mA3 mB2 WT mA3 mB2
endo-NPM1 + + + - - -
E.
Sucrose gradient, 10-30%
A260 (a.u.)
Fraction number
NPM1WT
NPM1mA3
NPM1mB2
NPM1WT-KO
NPM1mA3-KO
NPM1mB2-KO
F.
Overexpressed construct
Endogenous NPM1
Fractions
NPMWT NPMmA3 NPMmB2
WT
KO
Input 1+2 3 4+5 6+7 8+9 10+11
RFP
RFP*
NPM1
BOP1
NOP2
NOP53
uS3
EtBr

**Figure S10: Further characterizing NPM1 mutants.** (A) Western blot with NPM1 antibody in all NPM1 mutant cell lines used. Various species of NPM1 indicated. (B) Western blot with antibody that recognizes NPM1 for IP of NPM1 mutant constructs. Fusion protein (IP) and endogenous NPM1 bands indicated. Bottom, quantification across two replicates. Error bars are range, n = 2. (C) Western blot of NPM1 versus NPM1 knock out cell lines with antibody that recognizes NPM1. Below, representative images of NPM1 mutants in NPM1 knock out cells. Inverted LUT. Scale bar is 10 μm. (D) qPCR for rRNA precursors bound to NPM1 mutants. Primers indicated in schematic above. Fold change relative to IP of NPM1-WT, dotted line indicates 1 (no change). (E) Sucrose gradient fractionation of nuclear pre-ribosomal complexes isolated from HeLa cells transduced with $NPM1^{WT}$-mScarlet (blue curve), $NPM1^{mA3}$-mScarlet (orange curve), and $NPM1^{mB2}$-mScarlet (yellow curve). Cells in knock-out background shown as dotted line. (F) Western blot analysis of proteins from pooled sucrose gradient fractions, as shown in E. Ethidium bromide stained gels of RNA from pooled fractions shown below. For knockout cell background, RFP* is the same blot as RFP but scaled to better show mB2 signal. Several panels are reproduced from Fig. 5 for reference.

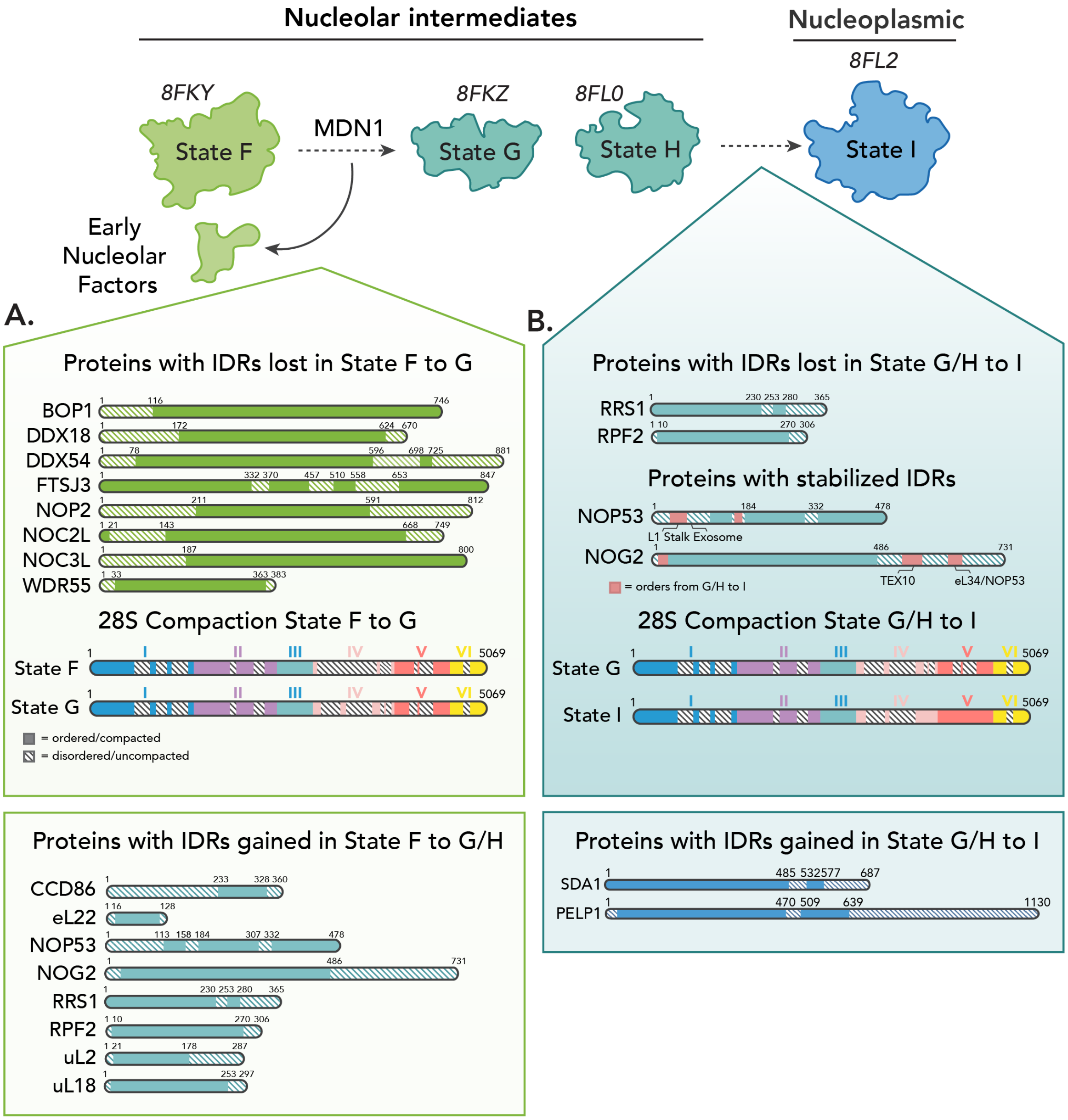


**Figure S11: Characterizing trans interactions in pre-LSU phenotypic clusters.** Top, schematic of pre-LSU intermediates associated with LSU1, LSU2, and nucleoplasmic export (States F, G/H, and I, respectively. PDB IDs indicated below). (A) Schematics for proteins with IDRS lost and gained at transition from LSU1 to LSU2 (State F to States G/H, respectively) and schematics of 28S showing uncompacted regions (unmodeled regions) for each state. (B) Schematics for proteins with IDRS lost and gained at transition from States G/H to State I and schematics of 28S showing uncompacted regions (unmodeled regions) for each state.
